# A central role for CCR2 in monocyte recruitment and blood-brain barrier disruption during Usutu virus encephalitis

**DOI:** 10.1101/2024.12.12.627941

**Authors:** Emily Slowikowski, Céleste Willems, Robertha Mariana Rodrigues Lemes, Sara Schuermans, Nele Berghmans, Rebeca Paiva Fróes Rocha, Erik Martens, Paul Proost, Leen Delang, Rafael Elias Marques, José Carlos Alves Filho, Pedro Elias Marques

## Abstract

Usutu virus (USUV) is an emerging neurotropic flavivirus capable of causing encephalitis in humans. Here, our main goal was to characterize the innate immune response in the brain during USUV encephalitis and to identify strategies to control disease severity. Using an immunocompetent mouse model of USUV encephalitis, we showed that microglia activation, blood-brain barrier (BBB) disruption and inflammatory monocyte recruitment are hallmarks of disease 6 days post infection. Activated microglia were in close association to USUV-infected cells, concomitantly with elevated levels of IL-6, IFN-γ, CCL2, CCL5, CXCL10 and CXCL1 in the brain. Monocyte recruitment was CCR2-dependent and driven by IFN-γ and CCL2 production beneath the brain vasculature. Moreover, CCR2 deficiency inhibited microglia activation and BBB disruption, showing the central role of CCR2 in USUV encephalitis. Accordingly, treatment with dexamethasone prevented pro-inflammatory mediator production and reduced leukocyte recruitment significantly, restraining encephalitis severity. Concluding, USUV encephalitis is driven by CCR2-mediated monocyte recruitment and BBB disruption, and blocked therapeutically by glucocorticoids.

**SUMMARY:** The neurotropic Usutu virus can cause encephalitis driven by CCR2-mediated monocyte recruitment, microglia activation and blood-brain barrier disruption, all of which are inhibited by glucocorticoid treatment.

## INTRODUCTION

Viral infections of the central nervous system (CNS) are a global public health threat, which are often caused by neurotropic viruses belonging to the *Flaviviridae* family. These vector-borne RNA viruses persist in enzootic cycles and are of particular concern due to their potential for (re)emergence due to environmental changes (Morens et al., 2004; Pierson & Diamond, 2020; Sips et al., 2012). Usutu virus (USUV) is an emerging flavivirus within the Japanese encephalitis virus serocomplex and is closely related to West Nile virus (WNV), with which it co-circulates in Europe. Since its introduction from Africa in Italy and Austria, USUV has rapidly spread across the European continent over the past two decades. Here, it has caused mass mortality events in birds, which act as amplifying hosts. Humans and other mammals can also be infected and develop disease (Vilibic-Cavlek et al., 2020). Importantly, seroprevalence studies in humans and animals have demonstrated active USUV circulation in both Europe and Africa, highlighting the ongoing threat of new USUV outbreaks (Angeloni et al., 2023; Tinto et al., 2022). A high degree of serological cross-reactivity between USUV and WNV complicates diagnostic discrimination and it is likely that many USUV cases have been misdiagnosed, underestimating the impact of USUV circulation (Llorente et al., 2019; Simonin, 2024). The majority of infections in humans are asymptomatic or cause febrile illness, nevertheless, severe neurological manifestations can develop in both healthy and immunocompromised individuals (Cadar & Simonin, 2022). Importantly, there are no approved antiviral drugs or human vaccines available for either USUV or WNV.

Viral encephalitis is a severe condition in which virus-induced brain inflammation gives rise to acute clinical manifestations such as headache, fever and altered consciousness. Neurological, cognitive and behavioural sequelae can persist until long after the virus has been cleared from the brain (John et al., 2015; Klein et al., 2019). Viral encephalitis is initiated when a neurotropic virus infects the brain, and it is detected by resident cells including microglia, astrocytes and neurons via pattern recognition receptors. Downstream signalling promotes the transcription of pro-inflammatory factors and induction of the type I interferon (IFN) response. This innate immune cascade leads to the release of cytokines and chemokines that recruit leukocytes from the periphery to infiltrate the perivascular space and parenchyma (Klein et al., 2017; McGavern & Kang, 2011). In general, monocytes and T cells are the most abundantly recruited cell populations, however, they can exert both protective antiviral functions and detrimental immunopathogenic effects. Another hallmark of CNS infection and the associated neuroinflammation is the disruption of the blood-brain barrier (BBB). Besides the additional facilitation of viral entry into the brain, vascular leakage can lead to severe clinical consequences, such as brain edema and increased intracranial pressure (Ashraf et al., 2021; Tyler, 2018). These events resonate in a complex interplay between viruses, cells and inflammatory mediators with discrepancies among different pathogens. Notably, USUV-induced encephalitis remains an almost unexplored field that has gained research interest in recent years. Abundant cytokine and chemokine expression has been detected in the brain of USUV-infected neonatal mice, together with the presence of T cells and macrophages (Clé et al., 2020). Even though chemokine-driven leukocyte infiltration is typical for viral encephalitis, the recruitment dynamics of monocytes to the brain (e.g. using intravital microscopy) has not yet been studied in USUV or any other flavivirus-induced encephalitis. Moreover, the mediators driving monocyte migration into the USUV-infected brain remain to be elucidated, as well as whether USUV compromises BBB integrity. USUV infection of *in vitro* BBB models and one *in vivo* model using IFNAR^−/−^ mice suggested that USUV does not affect BBB permeability directly (Clé et al., 2020; Constant et al., 2023). However, these studies do not exclude inflammation-induced BBB disruption.

Inflammatory monocytes play a complex role in neurological infections (Spiteri et al., 2022; Terry et al., 2012), of which the contribution of monocytes to USUV-induced encephalitis has yet to be explored. The majority of studies on monocytes in other neurotropic flavivirus infections have focused on their ability to be infected peripherally and carry the virus into the brain, a process known as the ‘trojan-horse’ mechanism (De Vries & Harding, 2023). Modulation of monocyte trafficking using neutralizing antibodies against CCL2 or VLA-4 integrin led to increased survival rates in an intranasal infection model of WNV (Getts et al., 2008, 2012, 2014). In contrast, a protective role for monocytes/macrophages during early WNV infection was observed, since viremia, neuroinvasion and mortality were increased in CCR2-deficient mice or treated with clodronate liposomes (Ben-Nathan et al., 1996; Bryan et al., 2018; Lim et al., 2011; Purtha et al., 2008; Winkelmann et al., 2014). These models are based on peripheral infections, leaving the role of monocytes during established brain infection and encephalitis still poorly understood. Likewise, existing mouse models of USUV infection have been conducted by peripheral inoculations in both wild-type (WT), immunocompromised or neonatal mice (Benzarti & Garigliany, 2020; Clé et al., 2020). However, these models fail to produce consistent neuroinvasive cases or result in a neurovirulent model that is not ideal for studying the immune response in the brain. An alternative is to inoculate adult immunocompetent mice via the intracranial route, achieving robust and consistent brain infection and inflammation.

Herein, we investigated the immune response triggered by USUV infection in the brain and demonstrated that monocytes are the most recruited leukocyte population. Using brain intravital microscopy, we investigated monocyte dynamics and vascular integrity of the BBB in both WT and transgenic (CCR2^−/−^ and IFN-γ^−/−^) mice. Furthermore, we tested different anti-inflammatory drugs to deepen our understanding of beneficial and detrimental aspects of the host’s immune response to neurotropic flavivirus infection.

## RESULTS

### USUV replicates in the brain and causes disease in immunocompetent mice

Given the limited susceptibility of immunocompetent mice to USUV infection, we inoculated mice via the intracranial route to ensure viral infection of the brain. By bypassing the skin infection, we obtained a consistent and effective model of encephalitis, reducing substantially the number of mice needed in this study. Adult C57BL/6J mice were injected with PBS as mock or USUV at 10^2^-10^4^ plaque forming units (PFU). Mice receiving 10^4^ PFU USUV, but not lower inocula lost a significant amount of body weight 6 days post infection (dpi) **(Fig. 1 A)**. USUV-infected mice (10^4^ PFU) presented general disease symptoms and at the end-stage typical signs of neurotropic infections such as paralysis of hind limbs. Moreover, several infected mice were affected by conjunctivitis of one or both eyes. USUV infection at a dose of 10^4^ PFU was lethal by day 7 dpi, however, mice infected with heat-inactivated USUV did not develop any signs of disease or mortality, showing that USUV-induced encephalitis requires local viral replication **(Fig. 1 B)**. Indeed, measurement of the infectious viral load in brain homogenates by end-point titration (TCID50) showed a dose-dependent increase corresponding to the initial inoculum **(Fig. 1 C)**. Considering that an inoculum of 10^4^ PFU induced a robust infection, all further experiments were conducted using this viral input.

**Figure 1.**
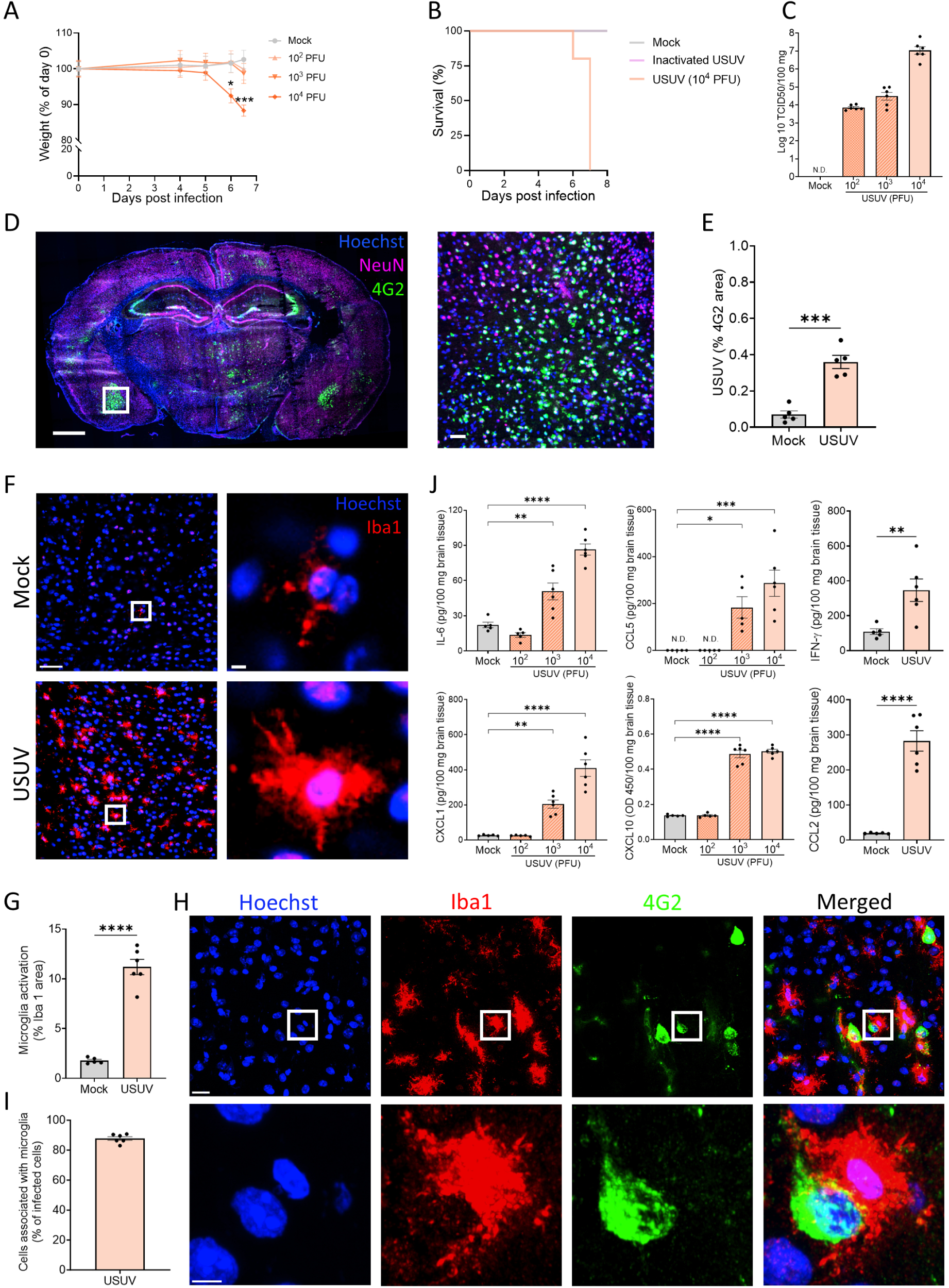
USUV infection in the brain induces acute encephalitis in immunocompetent mice. **(A)** Changes in body weight of mock or USUV-infected (10^2^-10^4^ PFU) mice. N=6 per group. **(B)** Survival of C57BL/6J mice inoculated with vehicle (mock), heat-inactivated or infective USUV. N=5 per group. **(C)** Infectious virus in the brain was measured at 6 dpi and is expressed as the log_10_-transformed 50% tissue culture infectious dose (TCID_50_) per 100 mg of brain tissue. **(D)** Confocal images showing immunostaining of brain cryosections from USUV-infected mice (10^4^ PFU) at 6 dpi, showing USUV infection (4G2, green), neurons (NeuN, magenta) and nuclei (Hoechst, blue). White square indicates the area zoomed. Scale bars = 1000 µm and 50 µm (zoom). **(E)** USUV infection is displayed as percentage of 4G2 stained area. **(F)** Representative images of microglia activation (Iba1, red) and nuclei (Hoechst, blue) in brain cryosections of mock and USUV-infected mice (10^4^ PFU) at 6 dpi. Scale bars = 50 µm and 5 µm (zoom). **(G)** Quantification of microglia activation, displayed as % of Iba1 stained area. **(H)** Representative images of immunostained USUV-infected cells (4G2, green) in the brain and microglia (Iba1, red). White squares indicate the zoomed areas. Scale bars = 20 μm and 5 μm. **(I)** Quantification of the percentage of infected cells that are associated with at least one microglial cell. **(J)** Levels of IL-6, CXCL1, CCL5, CXCL10, IFN-γ and CCL2 in full brain homogenates of mock and USUV-infected (10^2^-10^4^ PFU) mice at 6 dpi. Protein levels are displayed in pg per 100 mg of brain tissue or OD450 value (CXCL10). See also Fig S1 for TNF-α levels. Data are shown as mean ± SEM. N.D. = not detected. Image quantifications were pooled from minimum 5 pictures per mouse brain. Each dot in the graphs represents a single mouse. P values were obtained with two-way ANOVA (A), student’s t test (E, G, J) or one-way ANOVA followed by Dunnett’s multiple comparisons test (J). Significant differences compared to mock-infected mice are indicated with *P<0.05, **P<0.01, ***P<0.001, ****P<0.0001.

Productive USUV replication in the brain of immunocompetent mice was confirmed by immunostaining for USUV with the flavivirus group antigen antibody anti-4G2. Confocal images showed distinct areas of USUV infection with a significant increase in 4G2-stained area compared to mock-infected controls **(Fig. 1 D-E)**. USUV was detected in many brain regions, however, the images show infection of the hippocampus, as seen for Zika virus infections (Garber et al., 2019), and the amygdala. Images also indicate that USUV infects mainly neurons, however, sporadic microglia infections were observed **(Fig. S1 A)**. To determine whether USUV infection caused apoptosis of brain cells, we measured the level of cleaved caspase-3 in brain homogenates by ELISA. However, there was no elevation of this marker for apoptosis in comparison to mock-infected mice **(Fig. S1 B)**. Altogether, USUV has the ability to replicate in the brain and cause neurotropic disease in immunocompetent mice.

### USUV infection induces microglia activation and cytokine release

Knowing that USUV infects cells and replicates in the brain of mice, we aimed to assess the effect of USUV infection on resident immune cells. We performed immunostaining for Iba1 in brain cryosections of mice at 6 dpi to assess the status of microglia. USUV-infected brains showed numerous microglia with significantly increased Iba1 expression **(Fig. 1 F-G)**. Moreover, microglial cell morphology became more amoeboid in shape, with a larger cell body, thicker ramifications and frequent occurrence of multinucleation **(Fig. S1 C)** (Lee et al., 2024). Given that increased Iba1 is a recognized marker of microglial activation, these data showed that USUV infection triggered robust stimulation of microglia in the brain. Interestingly, activated microglia were often located in close proximity to USUV-infected cells, identified by 4G2 staining. In fact, over 85% of infected cells were associated with at least one microglial cell **(Fig. 1 H-I)**.

To further characterize the immune activation during USUV-induced encephalitis, we measured pro-inflammatory cytokine and chemokine levels in brain homogenates of mock and USUV-infected mice. IL-6, CCL5, CXCL1 and CXCL10 levels were increased in a dose-dependent manner to the inoculum. 10^4^ PFU of USUV also led to significantly higher IFN-γ and CCL2 levels in the brains of USUV-infected mice **(Fig. 1 J)**. Conversely, there was no difference in TNF-α levels, a mediator which is typically produced by activated microglia **(Fig. S1 D)**. These data suggest that USUV infection induces massive encephalitis in mice 6 dpi, a response marked by microglial activation and production of multiple pro-inflammatory cytokines and chemokines.

### USUV infection causes immunometabolic activation of macrophages

Considering the extensive activation of microglia by USUV, we hypothesized that the virus led to alterations in cellular metabolism. To investigate this, we assessed parameters of glycolysis in bone marrow-derived macrophages (BMDM) infected with USUV. BMDMs were also incubated with different comparative stimuli for 24h: medium as mock; LPS (100 ng/mL) as a proxy for bacterial stimulation; poly I:C (100 ng/mL) as representative of exposure to viral RNA; USUV at multiplicity of infection (MOI) 1 and 10. The extracellular acidification rate (ECAR) was measured in BMDMs first in glucose-free medium, showing ECAR levels that can be attributed to other sources than glycolysis such as the tricarboxylic acid (TCA) cycle. Glucose was added subsequently, causing an increased ECAR ascribed to increased glycolysis. Addition of oligomycin, an inhibitor of mitochondrial ATP synthase, pushes the cells to use glycolysis to its maximum capacity. Finally, the glycose analogue 2-deoxy-D-glucose (2-DG) acts as a competitive inhibitor and shuts down glycolysis, restoring the ECAR to non-glycolytic acidification **(Fig. 2 A)**. USUV at the highest MOI (10) caused a significant increase in glycolysis rate, glycolytic capacity and non-glycolytic acidification in comparison to mock, which were only partially elevated in BMDMs stimulated with LPS or poly I:C at the concentrations used, indicating that USUV infection is an immunometabolic modulator **(Fig. 2 B-D)**. Moreover, the extent of glycolytic alterations induced by USUV were once again inocula-dependent, connecting the intensity of leukocyte activation to the severity of viral infection. The glycolytic reserve was not affected by any of the stimuli **(Fig. S1 E).** The metabolic changes observed were also correlated to the level of inflammatory mediators secreted in the supernatant of USUV-infected BMDMs. Both CCL2 and CCL5 were significantly increased upon both LPS or USUV (MOI 10) incubation and increases in CCL2 were also measured after poly I:C or USUV (MOI 1) stimulation **(Fig. 2 E-F)**. Only LPS caused a significant increase in IL-6 compared to mock, suggesting that macrophages are not the main source of IL-6 upon USUV infection **(Fig. 2 G)**. Similarly, LPS but not USUV caused increased TNF-α levels, corroborating the lack of TNF-α production upon USUV infection *in vivo* **(Fig. S1 F)**.

**Figure 2.**
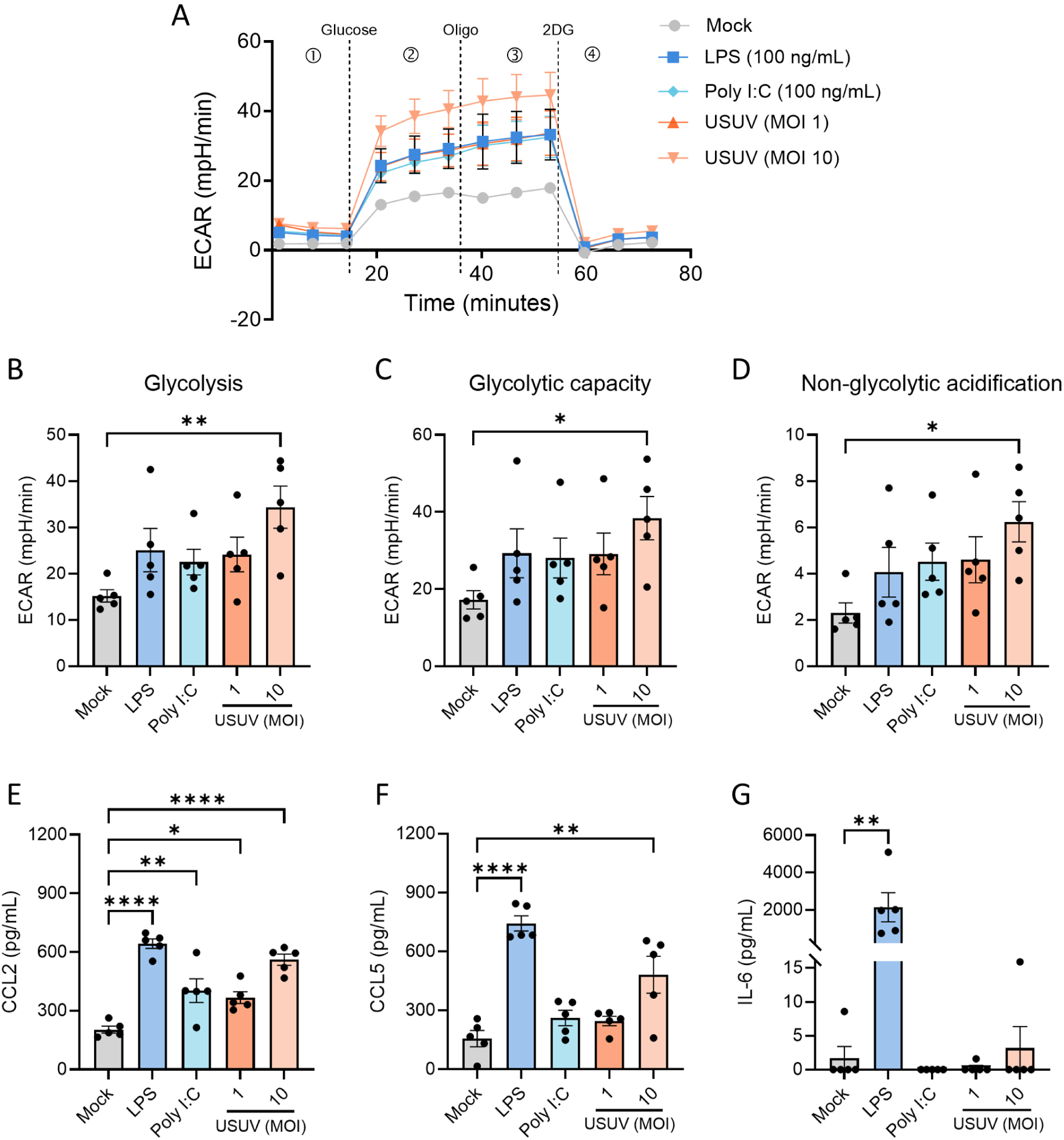
USUV increases glycolysis and cytokine release in bone marrow-derived macrophages. Bone marrow-derived macrophages (BMDM) were stimulated with medium (mock), LPS (100 ng/mL), poly I:C (100 ng/mL) or Usutu virus (USUV) at multiplicity of infection (MOI) 1 or 10 for 24h. **(A)** Kinetic extracellular acidification rate (ECAR) of BMDMs in response to glucose, oligomycin and 2-deoxyglucose (2-DG) measured by a Seahorse XF96 analyzer. **(B-D)** Glycolysis parameters calculated from ECAR values: glycolysis; glycolytic capacity; non-glycolytic acidification. **(E-G)** Levels of CCL2, CCL5 and IL-6 in supernatants of BMDMs after 24h of stimulation. Data are shown as mean ± SEM. Each dot in the graphs represents a single mouse. P values were obtained with one-way ANOVA followed by Dunnett’s multiple comparisons test. Significant differences compared to mock-infected mice are indicated with *P<0.05, **P<0.01, ***P<0.001, ****P<0.0001.

### Inflammatory monocytes are abundantly recruited to USUV-infected brains

Increased levels of the chemokines CCL2, CXCL1, CCL5 and CXCL10 induced by USUV infection **(Fig. 1 J)** are indicative of peripheral leukocyte recruitment to the brain. To evaluate this, we initially performed flow cytometry analysis of leukocytes in the brain of mock and USUV-infected mice. Under homeostatic conditions, the brain parenchyma maintains minimal leukocyte numbers (Török et al., 2021). Indeed, we found only a small fraction of CD45^hi^ leukocytes in mock-infected brains, however, leukocytes were massively recruited upon USUV infection **(Fig. 3 A)**. Inflammatory monocytes (CD45^hi^ Ly6C^hi^ CD11b^+^) were the most abundant cell type recruited to infected brains, followed by T cells (CD45^hi^ CD3^+^) **(Fig. 3 B-C and Fig. S1 G)**. Interestingly, the mean fluorescence intensity (MFI) of the integrin CD11b was increased on monocytes in infected brains, indicating that they become activated during encephalitis **(Fig. S1 H)**. Conversely, neutrophils, B cells and NK cells were not recruited following USUV infection **(Fig. 3 D-F)**. Monocyte recruitment was confirmed using confocal intravital microscopy (IVM) of the brain cortex. Prior to imaging, mice were injected intravenously with fluorescent dextran to enable visualization of the vasculature. Additionally, anti-CCR2 and anti-Ly6G antibodies were administered to label classical monocytes and neutrophils, respectively. Again, we stained the integrin CD11b as an activation marker for both cell types. Mock-infected mice showed solely the brain vasculature and occasional rolling leukocytes. In contrast, IVM imaging of USUV-infected brains exhibited an abundant recruitment of CCR2^+^ monocytes, many of which were either firmly adhered to or rolling/crawling along the vasculature **(Fig. 3 G and Video S1)**. In addition, monocytes were often expressing high levels of CD11b, confirming their activated and inflammatory state. Visualization of neutrophils was infrequent in both mock and USUV-infected brains, corroborating the flow cytometry results. These data show that leukocyte recruitment during USUV-induced encephalitis is extensive and predominantly composed by inflammatory CCR2^+^ monocytes.

**Figure 3.**
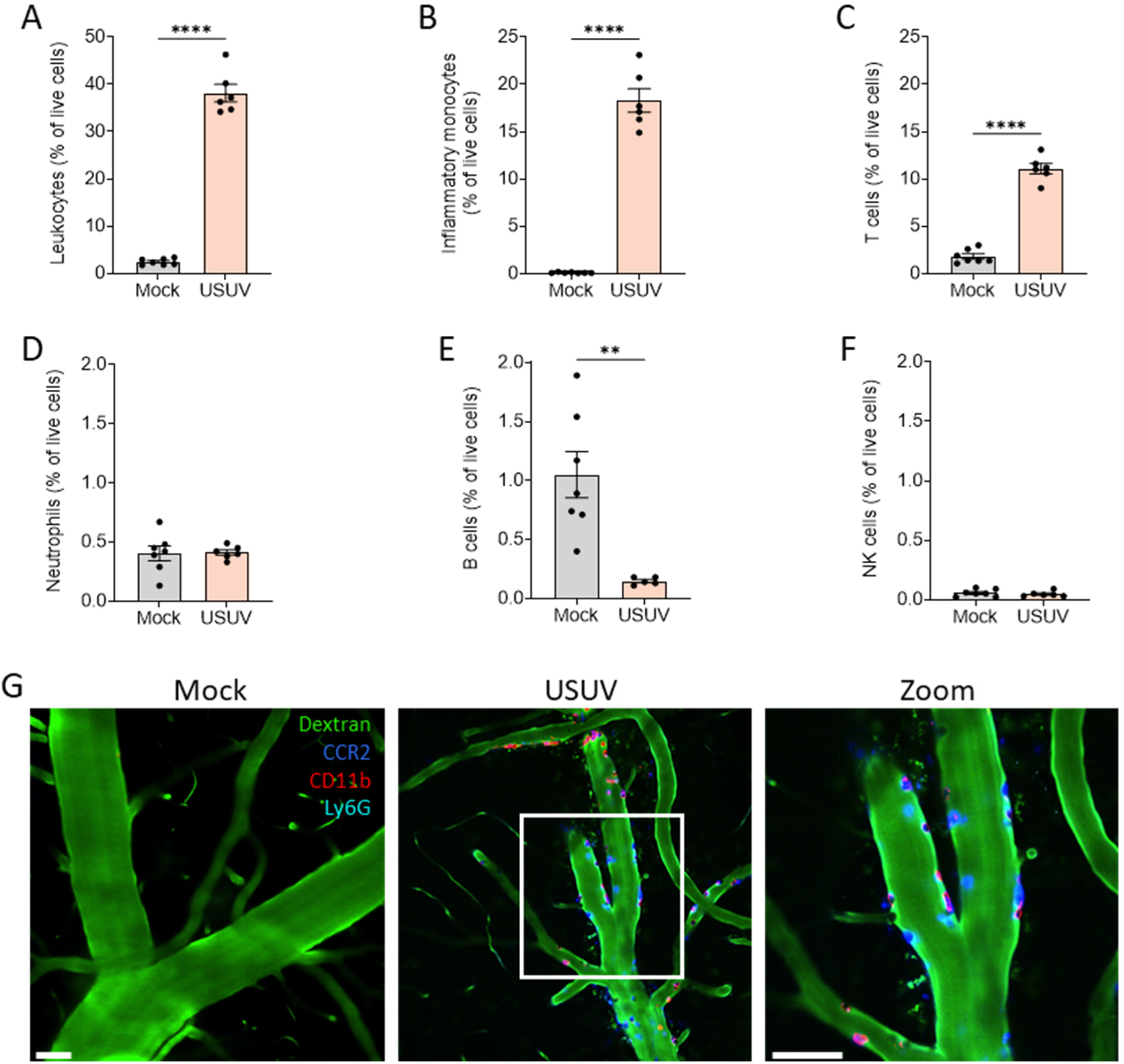
Inflammatory monocytes are abundantly recruited to the brain following USUV infection. Flow cytometry of cells isolated from the brain of mock and Usutu virus (USUV)-infected (10^4^ PFU) mice at 6 dpi. Percentages of **(A)** total leukocytes (CD45^high^), **(B)** inflammatory monocytes (CD45^high^, Ly6G^-^, CD3^-^, Ly6C^+^, CD11b^+^), **(C)** T cells (CD45^high^, CD3^+^), **(D)** neutrophils (CD45^high^, Ly6G^+^), **(E)** B cells (CD45^high^, CD19^+^) and **(F)** NK cells (CD45^high^, NK1.1^+^) within the live cell population are shown. **(G)** Representative confocal brain intravital microscopy (IVM) images (Video S1) showing the cortical region of mock and USUV-infected (10^4^ PFU) mice at 6 dpi. Brain vasculature (dextran, green), monocytes (CCR2, blue), neutrophils (Ly6G, cyan) and CD11b^+^ cells (CD11b, red) were visualized. White square indicates the zoomed area. Scale bars = 50 μm. Data are shown as mean ± SEM. Each dot in the graphs represents a single mouse. P values were obtained with student’s t test. Significant differences compared to mock-infected mice are indicated with *P<0.05, **P<0.01, ***P<0.001, ****P<0.0001.

### Monocyte recruitment to the brain depends on CCR2

Considering that CCR2 is a key chemokine receptor in classical monocytes and that we found elevated CCL2 levels in the brains of USUV-infected mice, we investigated the role of CCR2 in USUV-induced encephalitis. We observed that CCR2 is essential based on the significant reduction in total leukocytes recruited to the brain in CCR2^−/−^ mice **(Fig. 4 A)**. This was mainly a consequence of the substantial inhibition of monocyte and T cell infiltration in the brains of CCR2^−/−^ mice in comparison to WT. In contrast, the number of neutrophils was proportionally increased in CCR2^−/−^ mice, yet, still remaining low **(Fig. 4 A)**. The lack of B cell and NK cell recruitment remained the same for both WT and CCR2^−/−^ mice. Moreover, we observed that CCR2-deficient monocytes that were still recruited expressed lower CD11b levels, suggesting a less activated state and impairment in leukocyte adhesion **(Fig. 4 B)**. To further explore the role of CCR2 in monocyte recruitment to the brain, we performed brain IVM followed by monocyte tracking analysis **(Fig. 4 C-D)**. We found that monocytes lacking CCR2 were unable to adhere effectively to the brain vasculature, an integrin-dependent process crucial for brain infiltration **(Fig. 4 D)**. This led to a proportional increase in rolling cells, which travelled similar distances at normal speed compared to WT cells, indicating that selectin-mediated rolling is not impacted by CCR2 deficiency **(Fig. S2 A-B)**. Further investigation of CCR2^−/−^ mice showed that microglia were also less activated after USUV infection using Iba1 staining **(Fig. 4 E-F)**. Overall, our data indicate a dominant role of CCR2 in USUV-induced encephalitis, which drives the chemoattraction of monocytes and T cells, the effective adhesion of monocytes to the vasculature and full microglial activation.

**Figure 4.**
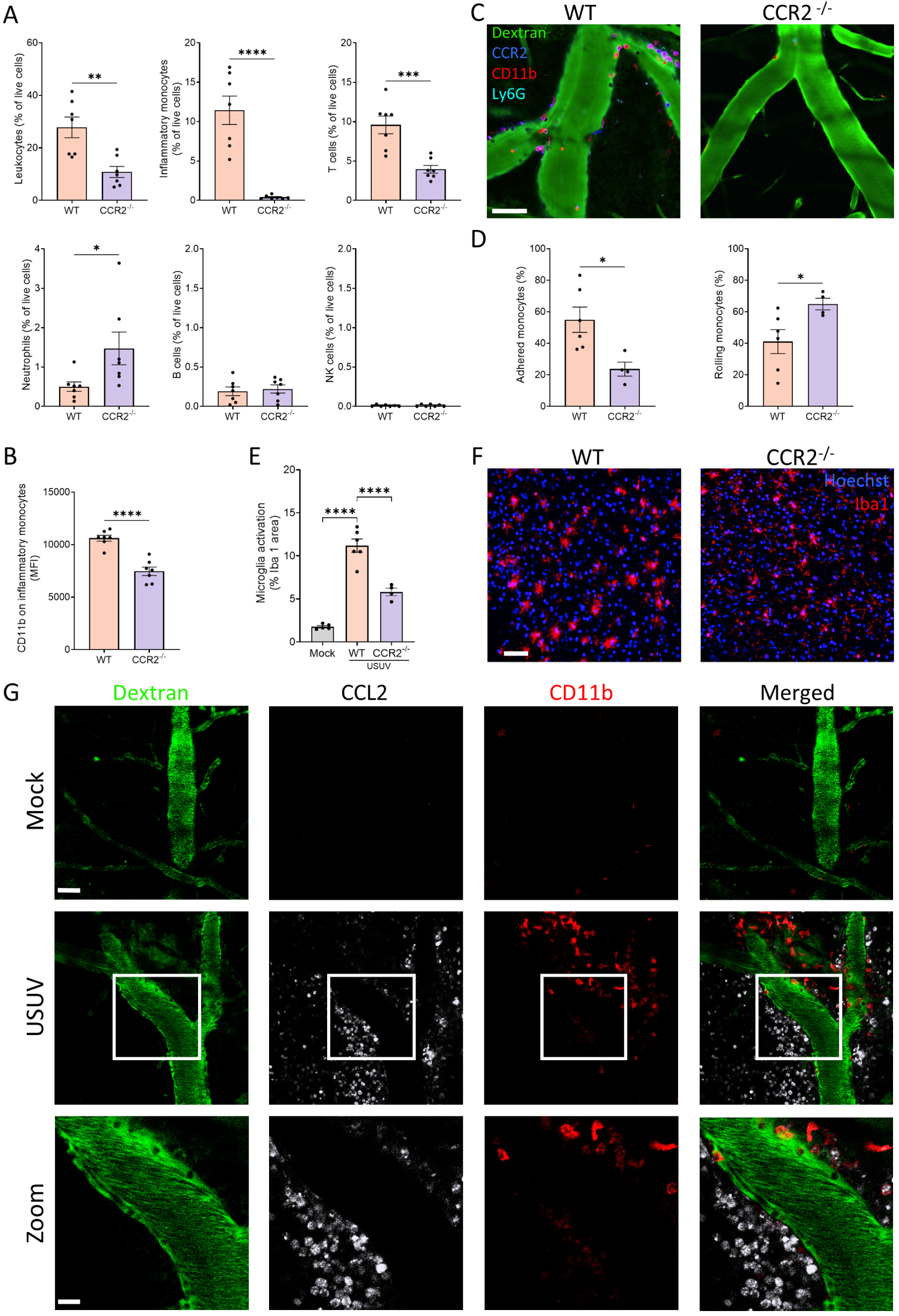
Inflammatory monocyte recruitment to the brain is CCR2 dependent. **(A)** Flow cytometry of cells isolated from the brain of wild-type (WT) and CCR2^−/−^ Usutu virus (USUV)-infected (10^4^ PFU) mice at 6 dpi. Percentages of total leukocytes (CD45^high^), inflammatory monocytes (CD45^high^, Ly6G^-^, CD3^-^, Ly6C^+^, CD11b^+^), T cells (CD45^high^, CD3^+^), neutrophils (CD45^high^, Ly6G^+^), B cells (CD45^high^, CD19^+^) and NK cells (CD45^high^, NK1.1^+^) within the live cell population are shown. **(B)** Mean fluorescence intensity (MFI) of CD11b on inflammatory monocytes by flow cytometry. **(C)** Representative confocal brain intravital microscopy images (Video S2) showing the cortical region of WT and CCR2^−/−^ USUV-infected (10^4^ PFU) mice at 6 dpi. Brain vasculature (dextran, green), monocytes (CCR2, blue), neutrophils (Ly6G, cyan) and CD11b^+^ cells (CD11b, red) were visualized. Scale bar = 50 μm. **(D)** Quantification of monocyte adherence state in the brain vasculature from intravital videos, displayed as the percentage of adhered and rolling monocytes. **(E)** Quantification of microglia activation in mock, WT and CCR2^−/−^ USUV-infected mice, displayed as percentage of Iba1 stained area. **(F)** Representative confocal images of immunostaining on brain cryosections, showing microglia (Iba1, red) and nuclei (Hoechst, blue). Scale bar= 50 μm. **(G)** Representative confocal brain intravital microscopy images from Ccl2-RFP^fl/fl^ mice inoculated with vehicle (mock) or USUV (10^4^ PFU) at 6 dpi. Brain vasculature (dextran, green), CCL2-producing cells (RFP, white) and CD11b^+^ cells (CD11b, red) were visualized. Scale bar = 50 μm. Data are shown as mean ± SEM. Each dot in the graphs represents a single mouse. P values were obtained with student’s t test (A, B, D) or one-way ANOVA followed by Dunnett’s multiple comparisons test (F). Significant differences compared to mock or WT infected mice are indicated with *P<0.05, **P<0.01, ***P<0.001, ****P<0.0001.

The levels of chemokines CCL2 and CXCL1 were elevated in CCR2^−/−^ brains following USUV infection **(Fig. S2 C-D)**, likely a compensatory mechanism from CCR2 deficiency. Conversely, pro-inflammatory cytokines IL-6, IFN-γ and the chemokine CXCL10 were not altered in CCR2^−/−^ mice **(Fig. S2 E-G)**. These data show that CCR2 is not required for the induction of pro-inflammatory mediator production in the infected brain. Furthermore, we observed that the survival rates of both CCR2^−/−^ and CCL2^−/−^ mice were comparable to those of WT mice **(Fig. S2 H)**, and that the viral load in brains of CCR2^−/−^ was also not altered **(Fig. S2 I)**. These data suggest that although CCR2 activation plays a central role in leukocyte recruitment during encephalitis, it is clearly a downstream event of the local immune response to USUV infection in the brain.

To broaden our understanding of CCL2 production and its spatial distribution, we performed brain IVM on USUV-infected CCL2-RFP^fl/fl^ mice, in which all cells expressing CCL2 concurrently express RFP. We observed at first that mock-infected brains had no detectable expression of CCL2 and negligible presence of CD11b-expressing cells, corroborating our previous data **(Fig. 4 G)**. In sharp contrast, USUV infection caused a massive increase in CCL2-expressing cells in the brain parenchyma, which were found both immediately below the endothelium and further into the brain tissue **(Fig. 4 G and Fig. S2 J)**, as identified by the blood vessels containing FITC-dextran. Moreover, the intravenous injection of anti-CD11b antibody showed a large population of stained cells in the proximity of brain blood vessels. Interestingly, given that the BBB is not permeable to antibodies and that the injection is performed minutes before imaging, this was suggestive of BBB disruption. Altogether, the production of CCL2 in USUV-infected brains is extensive and in close proximity to blood vessels, where it promotes CCR2-mediated leukocyte recruitment.

### CCR2^−/−^ mice are protected against BBB disruption during USUV encephalitis

Encephalitis involves multiple alterations in brain function, including disruption of the BBB and brain swelling. Therefore, we employed IVM to measure directly the vascular permeability of brain vessels using high molecular weight dextran (150 kDa). We observed that USUV infection induced increased vascular permeability in the brains of WT mice, which is consistent with the severity of viral encephalitis in these animals. Interestingly, mice deficient in CCR2 were protected, showing significantly less BBB disruption during USUV infection **(Fig. 5 A-B)**. These data confirm the induction of vascular leakage in USUV-infected brains and show the central role of CCR2 in this pathogenic process.

**Figure 5.**
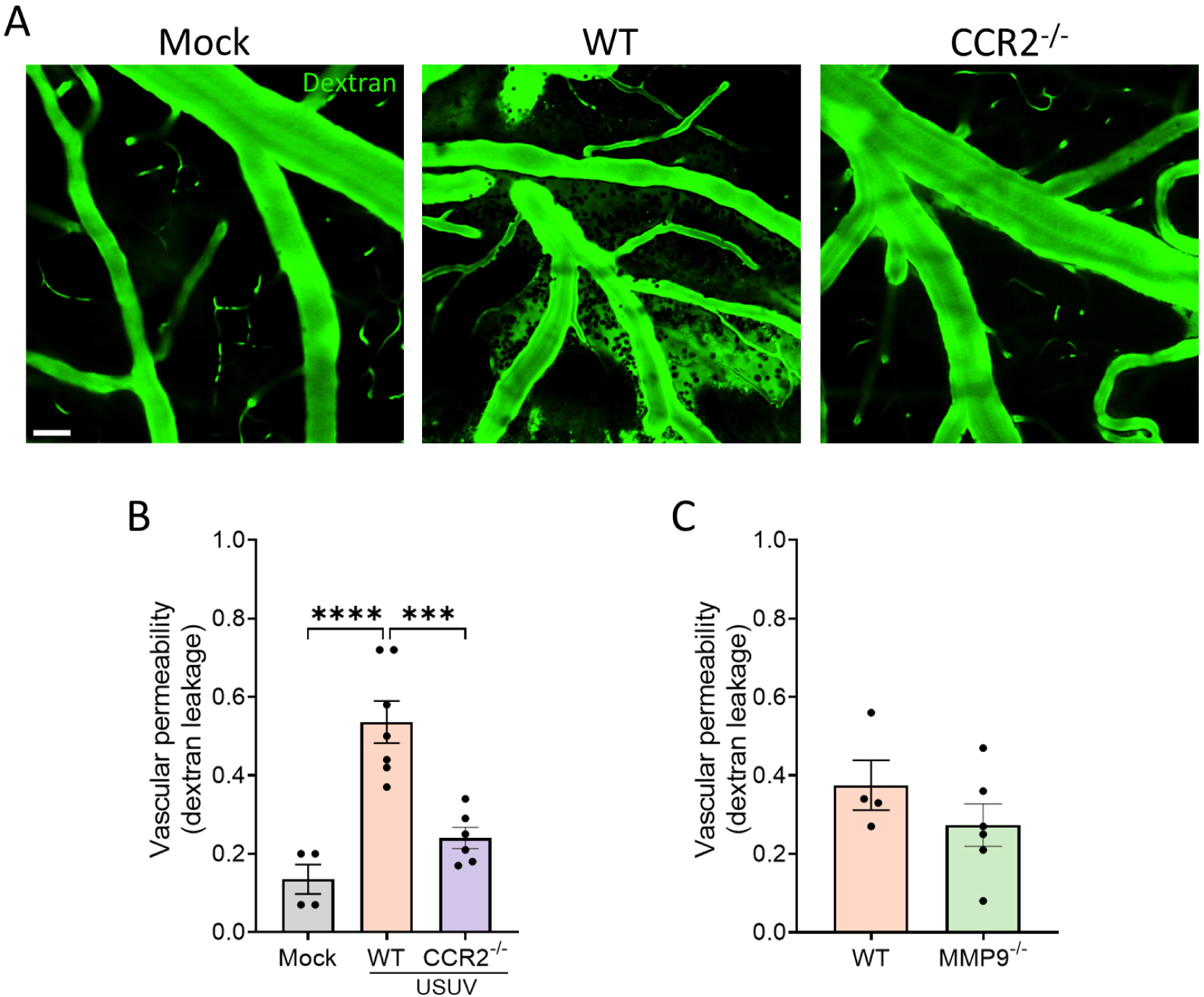
CCR2^−/−^ mice are protected against BBB disruption during USUV encephalitis. **(A)** Representative intravital images from mock, WT and CCR2^−/−^ USUV-infected mice, showing the brain vasculature and dextran leakage (dextran, green). Scale bar = 50 μm. Quantification of vascular permeability from cortical intravital images in **(B)** mock, WT, CCR2^−/−^ and **(C)** WT and MMP9^−/−^ USUV-infected mice. Dextran (150 kDa) leakage was measured by the ratio of dextran mean fluorescence intensity (MFI) outside versus inside the vessels. Data are shown as mean ± SEM. Each dot in the graphs represents a single mouse. P values were obtained with one-way ANOVA followed by Dunnett’s multiple comparisons test (B) or student’s t test (C). Significant differences compared to mock or WT infected mice are indicated with *P<0.05, **P<0.01, ***P<0.001, ****P<0.0001.

Matrix metalloproteases (MMPs) and in particular MMP9 are known to disrupt the BBB in multiple neuroinflammatory conditions (Vandooren et al., 2014), therefore, we measured MMP9 levels in the brain of mock, WT and CCR2^−/−^ USUV-infected mice using gelatine zymography **(Fig. S3 A-B)**. Pro-MMP9 levels were elevated in the brains of USUV-infected WT mice, yet, the elevation in pro-MMP9 was much more significant in CCR2^−/−^ and IFN-γ^−/−^ infected brains. When measuring the active form of MMP9, we observed that only CCR2^−/−^ mice had a significant elevation of MMP9 in the brain. We also measured the vascular leakage in the brain of USUV-infected MMP9^−/−^ mice and found it to be identical to WT mice **(Fig. 5 C)**, indicating that MMP9 activity is unrelated to the onset of BBB disruption in USUV encephalitis. Further characterization of USUV infection in MMP9-deficient mice showed no differences in weight loss **(Fig. S3 C)**, even though the viral load in the brain increased in MMP9^−/−^ mice **(Fig. S3 D)**. IL-6 production, but not CCL2, CXCL1 and CXCL10, was increased in MMP9^−/−^ mice compared to WT **(Fig. S3 E)**. Moreover, T cells, B cells, neutrophils and NK cells were more abundant in MMP9^−/−^ brains **(Fig. S3 F)**. Therefore, we conclude that MMP9 is not a key player in USUV-induced vascular leakage and that MMP9 deficiency is rather harmful to USUV-infected mice.

### IFN-γ triggers CCL2 production and inflammation in the brain

After establishing the role of the CCL2-CCR2 axis in USUV-induced encephalitis, we aimed to determine the stimuli triggering brain inflammation after USUV infection, including the production of CCL2. Considering the pro-inflammatory roles of IFN-γ upon neurotropic infections, including microglia activation (Kann et al., 2022), we investigated whether IFN-γ deficiency would impair the immune response to USUV. CCL2 levels measured in brain homogenates were significantly decreased in USUV-infected IFN-γ^−/−^ mice compared to WT mice **(Fig. 6 A)**. Also, IL-6, CXCL1 and CXCL10 were reduced in IFN-γ^−/−^ mice, indicating that IFN-γ drives the production of several pro-inflammatory mediators during USUV infection. Consequently, the knockout mice presented less inflammatory monocyte migration to the brain, alongside a minor inhibition in recruitment of other CD45^+^ leukocyte populations **(Fig. 6 B and Fig. S4 A)**. In addition, we employed IVM to inquire the parameters of monocyte recruitment in the brain vasculature of IFN-γ^−/−^ mice. Consistent with our previous data, monocytes in IFN-γ^−/−^ mice had impaired adhesion but increased rolling in the brain vasculature during USUV infection **(Fig. 6 C)**. Moreover, both microglia activation **(Fig. 6D)** and vascular permeability **(Fig. 6 E)** were reduced in infected IFN-γ^−/−^ mice. Again, we found no differences in body weight **(Fig. S4 B)** or viral load **(Fig. S4 C)** between WT and IFN-γ^−/−^ mice. It is interesting to note that the IFN-γ^−/−^ phenotype resembled that of CCR2^−/−^ mice although to a lesser extent, likely due to the levels of CCL2 and other inflammatory mediators not being completely reduced in IFN-γ^−/−^ mice. Importantly, IFN-γ^−/−^ mice presented lower CCL2 production, but CCR2^−/−^ mice had identical IFN-γ levels **(Fig. S2 F)**, suggesting that IFN-γ is an upstream inducer of CCL2 production. Overall, the data suggests that IFN-γ drives brain inflammation to its full extent during USUV infection, as evidenced by the dampening of pro-inflammatory mediators, monocyte recruitment, microglia activation and BBB disruption in IFN-γ^−/−^ mice.

**Figure 6.**
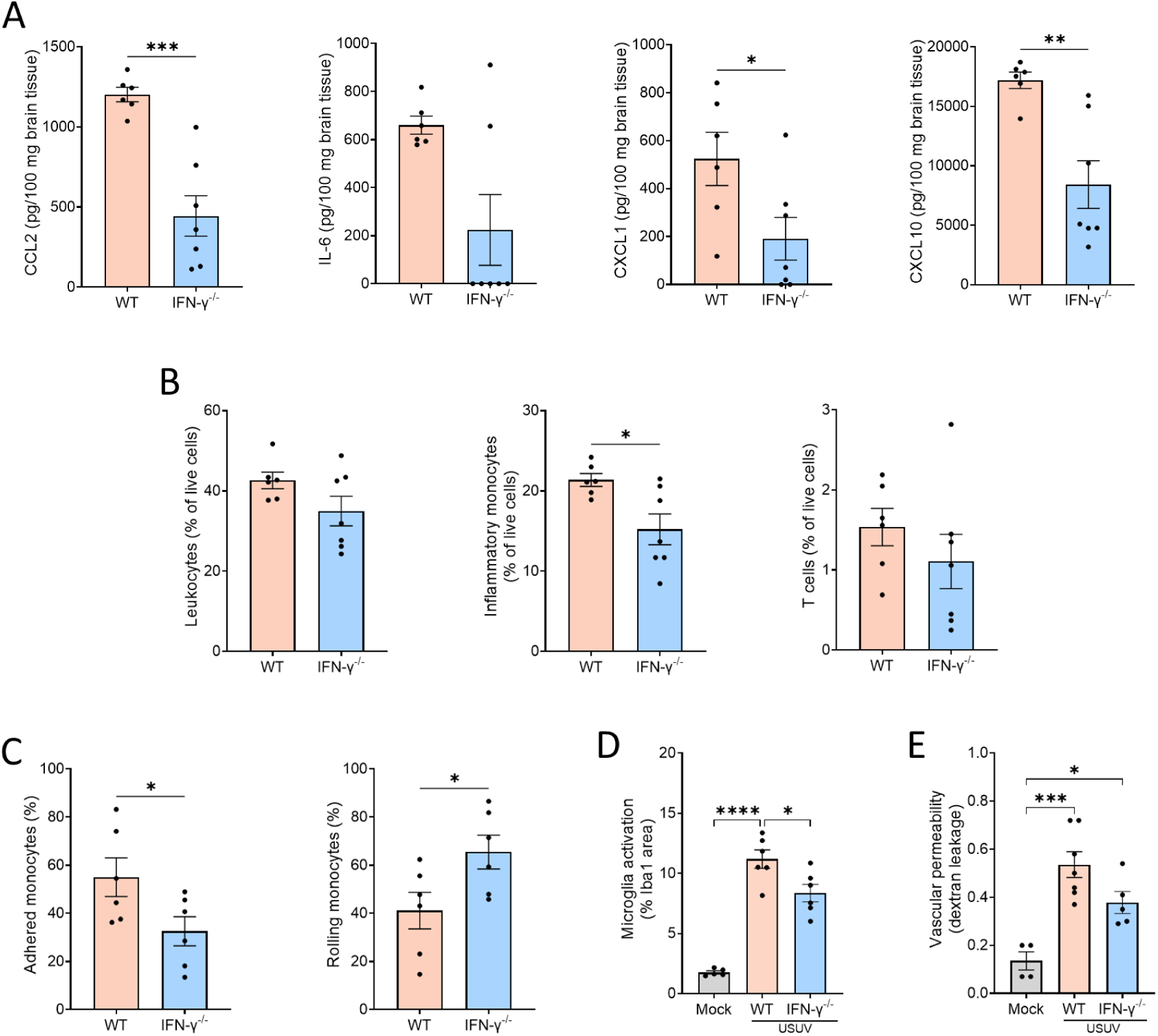
IFN-γ triggers CCL2 production and inflammation in the brain. **(A)** Cytokine levels of CCL2, IL-6, CXCL1 and CXCL10 in full brain homogenates of WT and IFN-γ^−/−^ USUV-infected (10^4^ PFU) mice at 6 dpi. Protein levels are displayed in pg per 100 mg of brain tissue. **(B)** Flow cytometry of cells isolated from the brain of WT and IFN-γ^−/−^ USUV-infected mice. Percentages of total leukocytes (CD45^high^), inflammatory monocytes (CD45^high^, Ly6G^-^, CD3^-^, Ly6C^+^, CD11b^+^) and T cells (CD45^high^, CD3^+^) within the live cell population are shown. **(C)** Quantification of monocyte adherence state in the brain vasculature from intravital videos, displayed as the percentage of adhered and rolling monocytes. **(D)** Quantification of microglia activation in mock, WT and IFN-γ^−/−^ USUV-infected mice, displayed as percentage of Iba1 stained area. Mock and WT groups are identical to Fig.4E. **(E)** Quantification of vascular permeability from cortical intravital images. Dextran (150 kDa) leakage was measured by the ratio of dextran mean fluorescence intensity (MFI) leaked outside the vessel over inside the vessel. Mock and WT groups are identical to Fig. 5B. Data are shown as mean ± SEM. Each dot in the graphs represents a single mouse. Image quantifications were pooled from minimum 5 pictures per mouse brain. P values were obtained with student’s t test (A-C) or one-way ANOVA followed by Dunnett’s multiple comparisons test (D-E). Significant differences compared to mock or WT infected mice are indicated with *P<0.05, **P<0.01, ***P<0.001, ****P<0.0001.

### Dexamethasone reduces brain inflammation and disease severity in USUV encephalitis

Considering the extent of the inflammatory response in USUV-infected brains, including cytokine and chemokine production, recruitment of leukocytes and disruption of the BBB, we evaluated whether different commonly used anti-inflammatory drugs would present therapeutic potential to reduce encephalitis severity. For this purpose, mice were injected subcutaneously with aspirin (100 mg/kg), dexamethasone (50 mg/kg), minocycline (40 mg/kg) or vehicle (PBS) at days 4, 5 and 6 post infection **(Fig. 7 A)**. The choice for these drugs relied on their distinct mechanisms of action. Aspirin is a non-steroidal anti-inflammatory drug (NSAID) acting through the inhibition of cyclooxygenases (Vane & Botting, 2003). Dexamethasone is a potent (steroidal) glucocorticoid and minocycline is a tetracycline with anti-inflammatory and neuroprotective properties (Garrido-Mesa et al., 2013). All treatment groups showed lower IL-6 and CCL2 levels in the brain compared to vehicle treated mice **(Fig. 7 B-C)**. However, only dexamethasone significantly reduced IFN-γ and CXCL10 **(Fig. 7 D-E)**. Similarly, dexamethasone was most efficient in inhibiting leukocyte recruitment to the brain, including specific reductions in recruitment of inflammatory monocytes, T cells and to a lesser extent neutrophils **(Fig. 7 F-I)**. In parallel, we observed that the most effective drug for inhibiting monocyte recruitment was minocycline **(Fig. 7 G)**. NK cell numbers were slightly increased in the aspirin and minocycline groups but remained very scarce, and there were no changes in B cell populations **(Fig. 7 J-K)**. Since dexamethasone was the most promising anti-inflammatory therapy, we decided to investigate this treatment further. Interestingly, dexamethasone was able to reduce disease symptoms such as loss of weight **(Fig. 7 L)**, and caused a significant reduction in brain vascular leakage **(Fig. 7 M)** and microglia activation **(Fig. 7 N)** in response to USUV infection. However, this broad inhibition of the immune response against USUV also led to a significantly higher viral load in brains of infected mice **(Fig. 7 O)**.

**Figure 7.**
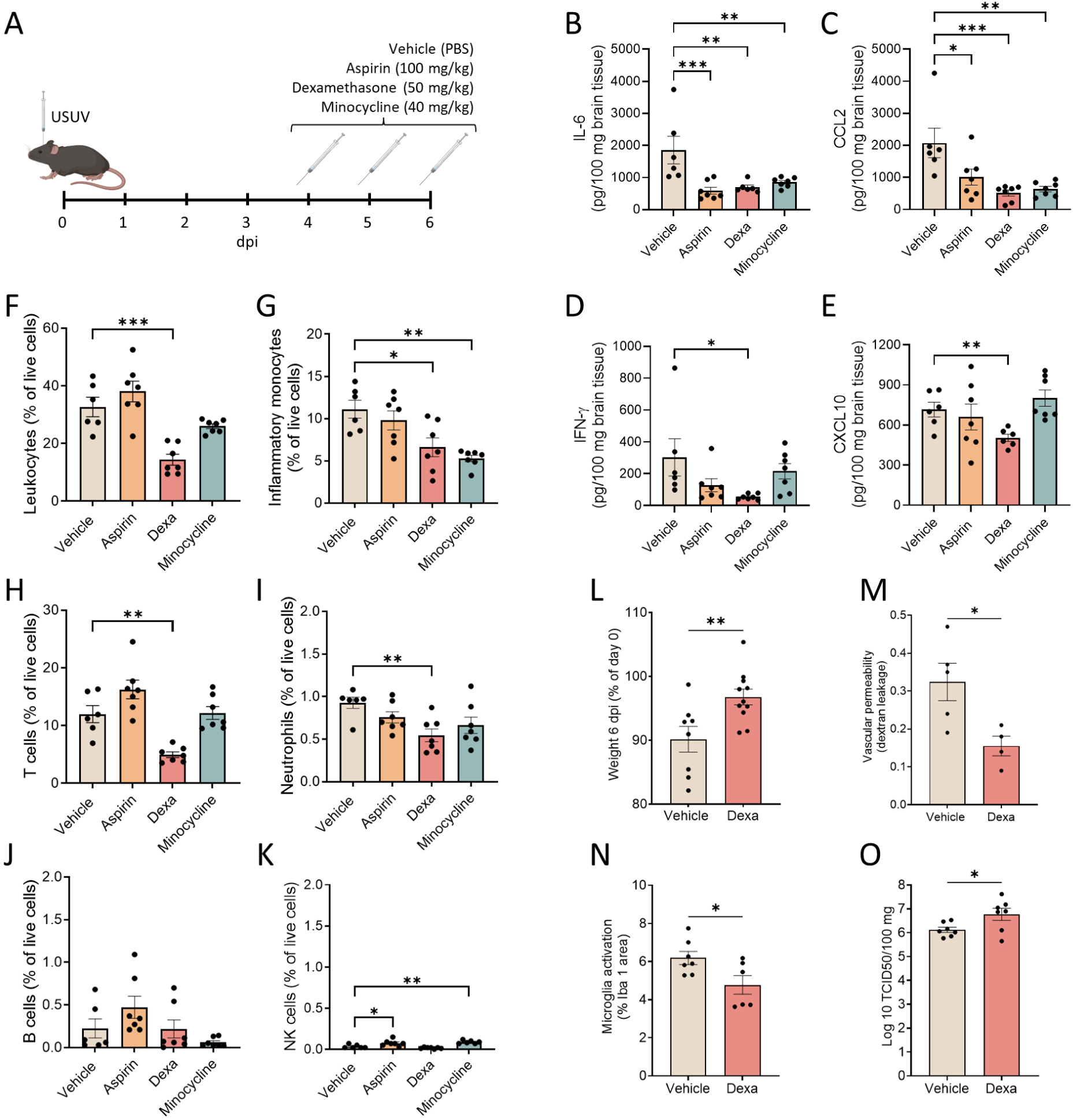
Dexamethasone reduces USUV-induced encephalitis and disease symptoms. **(A)** C57BL/6 mice were inoculated intracranially with USUV (10^4^ PFU) and treated subcutaneously with vehicle (PBS), aspirin (100 mg/kg), dexamethasone (50 mg/kg) or minocycline (40 mg/kg) at 4, 5 and 6 dpi. Cytokine and chemokine levels of **(B)** IL-6, **(C)** CCL2, **(D)** IFN-γ and **(E)** CXCL10 in brain homogenates at 6 dpi. Protein levels are displayed in pg per 100 mg of brain tissue. Flow cytometry of cells isolated from the brain of USUV-infected mice post-treatment. **(F)** Percentages of total leukocytes (CD45^high^), **(G)** inflammatory monocytes (CD45^high^, Ly6G^-^, CD3^-^, Ly6C^+^, CD11b^+^), **(H)** T cells (CD45^high^, CD3^+^), **(I)** neutrophils (CD45^high^, Ly6G^+^), **(J)** B cells (CD45^high^, CD19^+^) and **(K)** NK cells (CD45^high^, NK1.1^+^) within the live cell population are shown. **(L)** Body weight at 6 dpi of mice treated with vehicle or dexamethasone, shown as percentage of 0 dpi. **(M)** Quantification of vascular permeability from cortical intravital images. Dextran (150 kDa) leakage was measured by the ratio of dextran mean fluorescence intensity (MFI) leaked outside the vessel over inside the vessel. **(N)** Quantification of microglia activation in vehicle and dexamethasone treated USUV-infected mice, displayed as percentage of Iba1 stained area. **(O)** Infectious virus in the brain was measured at 6 dpi and is expressed as the log_10_-transformed 50% tissue culture infectious dose (TCID_50_) per 100 mg of tissue. Data are shown as mean ± SEM. Each dot in the graphs represents a single mouse. Image quantifications were pooled from minimum 5 pictures per mouse brain. P values were obtained with one-way ANOVA followed by Dunnett’s multiple comparisons test (B-K) or student’s t test (E, L-O). Significant differences compared to vehicle-treated mice are indicated with *P<0.05, **P<0.01, ***P<0.001, ****P<0.0001.

These data suggest that extensive inflammation rather than the viral load in the brain is at the root of USUV-induced disease burden. Indeed, we found a strong negative correlation between body weight loss and the mediators IFN-γ, CCL2 and CXCL10 **(Fig. S5 A)**. The correlation between IFN-γ and CCL2 is particularly strong with r > 0.8 **(Fig. S5 B)**, which corroborates our data that IFN-γ triggers CCL2 production **(Fig. 6 A)**. Altogether, dexamethasone treatment showed promising results in lowering the disease burden of USUV-induced encephalitis.

## DISCUSSION

USUV is an emerging flavivirus capable of neuroinvasion and causing CNS inflammation in humans (Cadar & Simonin, 2022). Despite its potential impact, the immune response in the brain upon USUV infection remains largely unexplored. In this study, we identified CCR2-driven leukocyte recruitment and BBB disruption as important mediators of USUV-induced immunopathology in the CNS. Consistent with other neurotropic infections, USUV triggered a strong activation of microglia. Notably, nearly all infected cells were found in close association with at least one microglial cell. While microglia are known to engulf dysfunctional and dying neurons (Prinz et al., 2019), we can only speculate that the observed microglia are phagocytosing the infected cells. Moreover, the production of inflammatory mediators by activated microglia and macrophages, including cytokines, reactive oxygen species (ROS) and MMPs, is an important contributor to neurodegeneration in CNS infections and neurodegenerative diseases (Kempermann & Neumann, 2003; Prinz et al., 2019). In the brain, pro-inflammatory cytokines such as IL-6 and IFN-γ were detected, whereas TNF-α was notably absent. While increased TNF-α expression in the murine brain during USUV infection has been reported, this study was conducted using neonatal Swiss mice, which may account for the discrepancy (Clé et al., 2021). Additionally, we detected the production of key chemokines CCL2, CXCL1, CCL5 and CXCL10, which play significant roles in recruiting immune cells to sites of infection.

Furthermore, we explored the impact of USUV infection on the metabolism of BMDMs and found a significant increase in glycolytic activity. The upregulation of glycolysis in macrophages is typically associated with a pro-inflammatory M1 phenotype, providing additional ATP to support antimicrobial functions such as phagocytosis and the production of inflammatory mediators (O’Neill et al., 2016; Russell et al., 2019). On the other hand, metabolic reprogramming is often exploited by viruses to enhance their replication, particularly in macrophages (Malemnganba et al., 2024; Russell et al., 2019). The glycolytic support for the replication of USUV has been shown in Vero cells (Wald et al., 2022). Thus, the enhanced glycolytic activity in macrophages during USUV infection may represent a double-edged sword, benefitting both host defence and viral propagation. Further research is necessary to fully understand this balance.

In the supernatants of infected BMDMs, we noted a significant increase in the chemokines CCL2 and CCL5, however, there was no corresponding rise in the pro-inflammatory cytokines IL-6 and TNF-α. This finding suggests that the source of IL-6 in the brain is likely of non-macrophage origin. This is further supported by *in vitro* experiments showing that microglia infected with USUV (EU3 strain) did not produce IL-6, whereas infected astrocytes did secrete the cytokine (Clé et al., 2021). Importantly, strain-dependent variations in cytokine response and virulence add complexity to understanding the immunopathology of USUV.

Leukocyte infiltration into the brain parenchyma is a hallmark of viral encephalitis. Similar to other flavivirus infections, USUV infection leads to the abundant chemokine-driven recruitment of monocytes and T cells into the brain (Glass et al., 2005; Lim et al., 2011; Michlmayr et al., 2014). Our findings demonstrate that inflammatory monocytes migrate to the brain in a CCR2-dependent manner, as these cells are almost entirely absent in CCR2-deficient mice. Moreover, the T cell population was largely reduced in CCR2-deficient mice, suggesting that T cells, which also express CCR2 (Luther & Cyster, 2001), are partially recruited via this receptor. Using intravital imaging, we show that rolling CCR2-deficient monocytes have difficulty to firmly adhere to the endothelium, which prevents them from undergoing diapedesis. This can be explained by the need for chemokine receptor signalling to stimulate surface expression and the inside-out activation of integrins, which is essential for interaction with adhesion molecules on endothelial cells mediating leukocyte arrest (Nourshargh & Alon, 2014).

It is still a matter of debate whether monocytes play a protective role in viral encephalitis by limiting viral replication or cause more harm in the process. Monocytes are generally regarded as protective in the early stage of flavivirus infections due to their role in peripheral viral clearance (Lim et al., 2011). Nevertheless, infected monocytes can contribute directly to viral dissemination by facilitating neuroinvasion via the ‘trojan-horse’ mechanism (De Vries & Harding, 2023). However, the use of the intracranial infection model limits our ability to investigate the neuroinvasion process and peripheral contribution of monocytes. Infiltration of inflammatory monocytes in the infected brain parenchyma and their differentiation can aid viral clearance through antigen presentation to T cells and release of pro-inflammatory and antiviral mediators. Nevertheless, both mechanisms can contribute to neuronal damage and BBB breakdown (Terry et al., 2012). In our study, we found that USUV-induced BBB leakage was strongly decreased in the absence of CCR2. A possible mechanism for reduced vascular leakage may involve the direct effect of CCR2 deficiency on endothelial cells. This hypothesis is supported by our observation that CCL2 was predominantly expressed by cells near the vasculature. CCL2-CCR2 signalling in brain endothelial cells has been shown to increase BBB permeability by remodelling of tight and adherens junction complexes (Roberts et al., 2012; Stamatovic et al., 2005). Moreover, CCR2 signalling promotes stress fiber formation resulting in cell retraction (Stamatovic et al., 2003). These changes can also result from signaling pathways induced by binding of leukocyte integrins to endothelial adhesion molecules (Cerutti & Ridley, 2017). As discussed earlier, chemokine receptor deficiency prevents the activation of leukocyte integrins. Consequently, the mere absence of massive leukocyte migration through the BBB in CCR2-deficient mice could itself alleviate associated vascular leakage. Furthermore, the strong reduction of monocytes in the brain may lead to a less inflammatory environment, potentially lowering the presence of mediators that contribute to BBB damage. The main cytokines that have been studied in this context include IL-6, TNF-α and IL-1β (Argaw et al., 2006; Rochfort & Cummins, 2015; Y. Wang et al., 2014). However, CCR2-deficient mice did not show decreased levels of measured cytokines or MMP9, suggesting that CCL2 is an end-point mediator in the neuroinflammatory process. Although MMP9 has been proposed to be a key mediator of WNV-induced BBB disruption (Verma et al., 2010; P. Wang et al., 2008), MMP9 does not appear to play a significant role in USUV-induced BBB leakage. Evidently, many other molecules secreted during neuroinflammation can influence BBB permeability, such as ROS, prostaglandins and vascular endothelial growth factor (VEGF). Therefore, the mechanism of BBB disruption needs further exploration in the context of USUV encephalitis.

Another interesting finding in CCR2-deficient mice was the lower activation of microglia in comparison to USUV-infected WT mice. This observation corroborates previous findings from various mouse models of both sterile and infectious neuroinflammation. For instance, mice infected with Japanese encephalitis virus (JEV) and treated with a CCR2 inhibitor exhibited significantly less microglial activation (Singh et al., 2020). Similarly, in a study of Theiler’s virus encephalitis, CCR2-deficient mice showed decreased microglial proliferation compared to infected WT mice (Käufer et al., 2018). Generally, microglia are thought to lack CCR2, indicating that the reduced microglial activation in CCR2-deficient mice is likely an indirect effect. This may be attributed to decreased monocyte infiltration into the brain, which limits crosstalk with microglia. Indeed, a study investigating microglia and macrophage phenotypes following traumatic brain injury showed alterations in the transcriptional profiles of microglia in CCR2-deficient mice. Recruited CCR2^+^ monocytes were associated with potentially neurotoxic subtypes of microglia promoting a type I IFN response (Somebang et al., 2021). In addition, inflammatory monocytes themselves can differentiate into a neurotoxic microglia-like phenotype, previously observed in WNV-encephalitis (Getts et al., 2008).

IFN-γ is a key player in driving cellular immunity during infectious diseases. While its actions are wide-ranging, it is particularly known for its function in activating and priming macrophages. Consequently, IFN-γ is an important regulator of microglia activation and inducer of pro-inflammatory cytokine release. Several studies have demonstrated the release of chemokines, including CCL2, by monocytes/macrophages and microglia upon IFN-γ stimulation (Boehm et al., 1997; Lin et al., 2009; Mcmanus et al., 2000; Rock et al., 2005). Since IFN-γ was abundantly expressed in the brain of USUV-infected mice, we investigated the impact of IFN-γ deficiency on CCL2 and cytokine release, microglia activation, leukocyte recruitment and BBB disruption. The phenotype of USUV-infected IFN-γ-deficient mice followed that of mice deficient in CCR2, yet to a lesser extent. Importantly, CCL2 levels in the brain were significantly lower in the absence of IFN-γ, corroborating above-mentioned studies on IFN-γ-dependent CCL2 induction. We suggest that the reduced leukocyte recruitment and vascular leakage observed in IFN-γ-deficient mice is at least partially attributed to the lower levels of CCL2.

Our findings point towards a mostly detrimental role of the innate immune response in USUV encephalitic disease. To substantiate this, we evaluated the effects of several approved anti-inflammatory drugs with distinct mechanisms of action. Treatment of USUV-infected mice with dexamethasone significantly reduced inflammation regarding cytokine production, leukocyte recruitment and microglia activation. Even though the viral load was higher in treated mice, BBB disruption and weight loss were significantly lessened, confirming that the host immune response rather than further viral replication is the main cause of USUV-induced disease. The use of corticosteroids such as dexamethasone for the treatment of WNV encephalitis has previously been proposed, however, case reports and clinical studies show conflicting results (Colaneri et al., 2023; Kal et al., 2022; Leis & Sinclair, 2019). Considering all this, it may be a useful approach to combine the use of steroids and antiviral drugs in USUV– and other flaviviral encephalitides.

## METHODS

### Animals

C57BL/6J mice were purchased from Janvier Labs. CCR2^−/−^, IFNg^−/−^ were obtained from the Jackson Laboratory. MMP9^−/−^ and its paired WT controls were provided by Prof. G Opdenakker. Mice were housed per gender in individually ventilated cages (maximum 5 mice per cage) at 21 °C and 12:12 light/dark cycles at the Animal facility of the Rega Institute (KU Leuven). B6.Cg-*Ccl2^tm1.1Pame^*/J (Ccl2-RFP^flox^) mice were purchased from The Jackson Laboratory and maintained in the Animal facility of FMRP-USP. Water and food were provided ad libitum and enrichment (bedding, toys and small houses) was provided. All animals used in this study were age 8-12 weeks. Both male and female mice were equally distributed across experiments, as no phenotypical differences were observed between genders. Housing conditions and experimental procedures were approved and performed following the guidelines of the Animal Ethics Committee from KU Leuven (registry number: P084/2022).

### Cells

Vero E6 cells (African green monkey kidney epithelial cells; ATCC: CRL-1586) were cultured in Dulbecco’s Modified Eagle Medium (DMEM, Lonza Biologics) supplemented with 10% fetal bovine serum (FBS, Sigma-Aldrich). Cells were incubated at 37°C and in the presence of 5% CO_2_.

### Mouse model of USUV infection

USUV strain SE/17 Europe 3 lineage (MK230892) was obtained from Prof. Mutien Garigliany (ULiège). A virus stock was produced on Vero E6 cells and the virus titer was determined by performing a plaque assay (Soto et al., 2023). Mice were inoculated intracranially with 1 x 10^4^ plaque-forming units (PFU) of USUV diluted in PBS or PBS only as mock-infected control, as based on the protocol created by Prof. Rafael Elias Marques (CNPEM, Brazil)(manuscript under review). Injections with a volume of 20 µL were performed using a 30G syringe (BD Micro-Fine) under isoflurane anesthesia (Abbot Laboratories Ltd.). The needle was inserted perpendicularly into the cranial cavity on the intersection of the medial and sagittal planes. At day 6 post-infection, mice were euthanized by exsanguination under anesthesia by ketamine (80 mg/kg) and xylazine (4 mg/kg). Tissue samples were used immediately or stored at −80°C until further analysis.

### Quantification of viral load

Collected hemi-brains were weighed, homogenized in 500 µL unsupplemented DMEM and centrifuged for 20 minutes (1000x g, 4 °C). Brain supernatants were used for end-point titration. Vero E6 cells were seeded in a 96-well plate and grown to confluence overnight. The next day, samples were added in serial dilutions and incubated at 37°C. After 5 days, the plates were checked for cytopathic effect and the viral titer was determined by using the Reed and Muench formula. Titers were expressed as Log10 50% tissue culture infective dose per 100 mg of brain tissue (Log10 TCID50/100 mg).

### Quantification of cytokines and chemokines

Concentrations of IL-6, IFN-γ, TNF-α, CCL2, CXCL1, CCL5, CXCL10 and cleaved caspase 3 were measured in cell supernatants and brain tissues by enzyme-linked immunosorbent assay (ELISA). Collected hemi-brains were weighed and homogenized in protein extraction buffer, consisting of 15 mM NaCl (VWR Chemicals, Radnor), 0.05% Tween 20 (Sigma-Aldrich), 5% Bovine Serum Albumin (BSA, Sigma-Aldrich), 10 mM EDTA (Sigma-Aldrich) and 1% protease inhibitor cocktail (Sigma-Aldrich) in PBS. Homogenates were centrifuged for 20 minutes (1000x g, 4 °C), supernatants were collected and proteins levels were quantified by sandwich ELISA (DuoSet R&D Systems) according to the manufacturer’s protocol. The absorbance was determined using a spectrophotometer (BioTek) at 450 nm. Results are expressed as pg per 100 mg of brain tissue or pg per mL of cell supernatant.

### Macrophage differentiation and stimulation

Bone marrow cells were aspirated from mice’s (C57BL/6J) dissected femur and cultured for 7 days in RPMI 1640 (Thermo Fisher Scientific) supplemented with FBS (20%), L929 conditioned medium (macrophage colony stimulating factors (M-CSF); 20%), GlutaMax (2 mM; Sigma Aldrich), and antibiotic-antimycotic (penicillin, streptomycin and fungizone; Thermo Fisher Scientific). Upon differentiation, BMDMs were seeded in 96-well plates (Seahorse XF96; Agilent Technologies) at 7,5 x 10^4^ cells/well (glycolytic function) or in 24-well plates at 3,75 x 10^5^ cells/well (cytokine quantification).

After adhesion, BMDMs were infected with USUV (Multiplicity Of Infection (MOI) 1 or 10) for 1 hour. Next, the supernatant was removed, cells were washed with PBS and cultured in RPMI 1640 supplemented with FBS (10%) and GlutaMax (2mM) for 24 hours. In parallel, positive control conditions were generated by stimulating BMDMs with LPS (100 ng/mL) or Poly I:C (100 ng/mL). All incubations were performed at 37 °C and in the presence of 5% CO_2_. Finally, cells were submitted to glycolytic function analysis in a Seahorse assay and the supernatants were harvested for cytokine and chemokine quantification.

### Glycolytic function

The extracellular Acidification Rate (ECAR) was measured in real-time using the Seahorse XF96 Analyser (Agilent) following the manufacturer’s instructions. Stimulated BMDMs were washed with XF media (non-buffered RPMI-1640 containing 2 mM L-glutamine and 1 mM sodium pyruvate; Agilent Seahorse XF24) and incubated for 1 hour at 37°C, in the absence of CO_2_. ECAR was measured under basal conditions and after adding the following to the cartridge of the Seahorse XF96 Analyzer: glucose (10 mM, Sigma Aldrich), oligomycin (1 μM; Sigma Aldrich) and 2-Deoxy-D-glucose (50 mM; Sigma Aldrich). The data were analyzed using Seahorse Wave Pro software.

### Cryosectioning and immunostaining

Brain tissues were carefully dissected, embedded in Polyfreeze (Sigma) and snap-frozen in liquid nitrogen. Brain cryosections of 14 μm thickness were cut using a cryostat (Microm Cryo-Star HM560, Thermo Fisher Scientific) and collected onto polysine slides (Epredia). The samples were fixed for 1 hour using 4% paraformaldehyde (PFA, Thermo Fisher Scientific) in PBS. The sections were washed with PBS and permeabilized using 0.1% Triton X-100 (Alfa Aesar) for 1 hour. The sections were washed again and blocked using 1% Fc block (Miltenyi Biotec) and 10% fetal bovine serum (FBS) for 1 hour. After another washing step, primary antibodies were added and incubated overnight (4°C). We stained the sections with rabbit anti-Iba1 (Abcam), mouse or rabbit anti-flavivirus group antigen 4G2 (Novus Biologicals) and mouse anti-NeuN (Abcam). The following day, slides were washed and secondary antibodies (10 μg/mL, Jackson ImmunoResearch) were added for 3 hours at room temperature. Specifications of all used antibodies can be found in supplementary **(Table S1)**. Nuclei were counterstained with Hoechst 33342 (10 μg/mL, Thermo Fisher Scientific), washed and mounting medium was applied (ProLong Diamond, Thermo Fisher Scientific). The slides were imaged using an Andor Dragonfly 200 spinning-disk confocal microscope (Oxford Instruments) equipped with a 25X objective, and analyzed using FIJI v1.54f and Imaris 9.9.1 software.

### Gelatin zymography

MMP9 from brain homogenates was purified using gelatin-Sepharose beads (GE Healthcare) on mini-spin columns (Biorad). Proteases were eluted from the beads with nonreducing loading buffer and run in 7.5% polyacrylamide gels containing 0.1% gelatin for electrophoretic protein separation. Refolding of the proteases was performed by washing the gels in a 2.5% Triton-X-100 solution. Next, the gels were incubated overnight in 10 mM CaCl_2_ and 50 mM Tris–HCl (pH 7.5, 37°C). Finally, the gels were stained with 0.1% Coomassie Brilliant Blue R-350 (GE Healthcare) and zymograms were analyzed using the ImageQuant TL software (GE Healthcare) (Vandooren et al., 2013).

### Cell isolation and flow cytometry

Dissected brain tissues were collected in cold RPMI 1640 medium (Biowest), manually minced and enzymatically digested by collagenase (0,5 mg/mL, Roche) for 30 minutes (37°C) on a shaker. Next, the tissue was strained through a 100 µm nylon cell strainer (pluriSelect) and centrifuged for 10 minutes (300x g, 4°C). The pellet was resuspended in 30% Percoll (Cytiva), added on top of 70% Percoll and gradient centrifuged for 30 minutes (500x g, RT) to separate immune cells from myelin debris. Cells were collected and washed twice with PBS and the pellet was resuspended in FACS buffer (PBS supplemented with 0.5% BSA and 2mM EDTA). The cell viability dye Zombie Aqua (Biolegend) and 1% Fc block (Miltenyi Biotec) in FACS buffer were added for 15 minutes (4°C). The samples were washed with FACS buffer and stained with different antibodies in brilliant stain buffer (BD Biosciences) for 25 minutes in the dark (4°C). After labeling, the cells were washed with FACS buffer, centrifuged for 3 minutes (300x g, 4°C) and resuspended in FACS buffer. To obtain data, samples were read in a Fortessa X20 (BD Biosciences) and analyzed using FlowJo 10.8.0 (FlowJo LLC) software. Antibody specifications and gating strategy can be found in the supplementary **(Table S1 and Fig. S6)**.

### Brain intravital microscopy

Mice were deeply anesthetized by subcutaneous administration of ketamine (80 mg/kg) and xylazine (4 mg/kg). A mixture of 150 kDa dextran-FITC (750 µg, Sigma-Aldrich) and conjugated antibodies (200 µg/kg) anti-mouse CCR2-BV421 (BioLegend), anti-mouse CD11b-PE (BD Biosciences) and anti-mouse Ly6G-Alexa Fluor 647 (BioLegend) diluted in PBS was injected intravenously **(Table S1)**. The mouse was placed under a stereoscopic microscope (Wild Heerbrugg) and hypothermia was prevented using a heat lamp (Beurer). Mineral oil (Sigma-Aldrich) was applied before the skin on top of the mouse head was removed. Next, a craniotomy was performed using a dental drill (Shiyang) with a drill bit (H1SE 204 014, Komet) to drill a square with a diagonal of approximately 5 mm in the parietal bone. Once the square bone was loose, PBS was added topically to ease its removal, carefully using fine tweezers. The mouse was positioned on a custom-made microscopy stage on which a drop of PBS prevents the exposed brain from drying out. The stage was placed in a heating unit at 37°C and the brain was imaged every 30 seconds for 20 minutes using a 25X objective in an Andor Dragonfly 200 series high speed 665 confocal platform system.

### Image analysis and monocyte tracking

Vascular leakage analysis was performed in FIJI v1.54f, by measuring the mean gray value of dextran-FITC signal inside the vessels and within the distance of 20 µm surrounding the vessels. Vascular permeability was determined as the ratio of dextran signals in and outside the vessels. Data were averaged from 5 different areas per mouse. Image alignment, monocyte tracking and morphology analysis were performed using the plugins Linear Stack Alignment with SIFT and TrackMate v7, in FIJI, following the developer’s methodology (Ershov et al., 2022). Proportions of adhered and rolling monocytes were counted manually. Leakage analysis and cell tracking was performed using at least 4 different animals per group.

### Statistical analysis

All data were analyzed using GraphPad Prism v9.3.1. and are represented as mean ± standard error of the mean (SEM). Normality was checked using the Shapiro-Wilkinson test and outliers were identified using the Grubb’s test. Normally distributed data were analyzed by a student’s t test, One-way or Two-way ANOVA. Non-parametric data were analyzed using the Mann-Whitney test or Kruskal-Wallis test. A p-value equal or lower than 0.05 was considered significant.

## Supporting information

Graphical abstract

## Acknowledgments

This work is supported by FWO-Vlaanderen Junior Research Grants (G058421N and G025923N), a KU Leuven C1 grant (C14/23/143), an FWO-FAPESP grant (G0F8822N) and the Rega Foundation. SS holds a PhD fellowship from FWO-Vlaanderen (1116922N). The intravital platform used in this research was obtained through the FWO medium-scale infrastructure funding (I009020N). This work was also supported by a FAPESP grant (2013/08216-2, Center for Research in Inflammatory Diseases) and the National Council for Scientific and Technological Development (CNPq – Brazil). REM is a CNPq research fellow and funded by INCT-ONE.

## Notes

### Competing Interest Statement

The authors have declared no competing interest.

## REFERENCES

1. Angeloni, G., Bertola, M., Lazzaro, E., Morini, M., Masi, G., Sinigaglia, A., Trevisan, M., Gossner, C. M., Haussig, J. M., Bakonyi, T., Capelli, G., & Barzon, L. (2023). Epidemiology, surveillance and diagnosis of Usutu virus infection in the EU/EEA, 2012 to 2021. Eurosurveillance, 28(33). 10.2807/1560-7917.ES.2023.28.33.2200929

2. Argaw, A. T., Zhang, Y., Snyder, B. J., Zhao, M.-L., Kopp, N., Lee, S. C., Raine, C. S., Brosnan, C. F., & John, G. R. (2006). IL-1β Regulates Blood-Brain Barrier Permeability via Reactivation of the Hypoxia-Angiogenesis Program. The Journal of Immunology, 177(8), 5574–5584. 10.4049/jimmunol.177.8.5574

3. Ashraf, U., Ding, Z., Deng, S., Ye, J., Cao, S., & Chen, Z. (2021). Pathogenicity and virulence of Japanese encephalitis virus: Neuroinflammation and neuronal cell damage. Virulence, 12(1), 968–980. 10.1080/21505594.2021.1899674

4. Ben-Nathan, D., Huitinga, I., Lustig, S., Van Rooijen, N., & Kobiler, D. (1996). West Nile virus neuroinvasion and encephalitis induced by macrophage depletion in mice. Archives of Virology, 141(3-4), 459–469. 10.1007/BF01718310

5. Benzarti, E., & Garigliany, M. (2020). In Vitro and In Vivo Models to Study the Zoonotic Mosquito-Borne Usutu Virus. Viruses, 12(10), 1116. 10.3390/v12101116

6. Boehm, U., Klamp, T., Groot, M., & Howard, J. C. (1997). CELLULAR RESPONSES TO INTERFERON-γ. Annual Review of Immunology, 15(1), 749–795. 10.1146/annurev.immunol.15.1.749

7. Bryan, M. A., Giordano, D., Draves, K. E., Green, R., Gale, M., & Clark, E. A. (2018). Splenic macrophages are required for protective innate immunity against West Nile virus. PLOS ONE, 13(2), e0191690. 10.1371/journal.pone.0191690

8. Cadar, D., & Simonin, Y. (2022). Human Usutu Virus Infections in Europe: A New Risk on Horizon? Viruses, 15(1), 77. 10.3390/v15010077

9. Cerutti, C., & Ridley, A. J. (2017). Endothelial cell-cell adhesion and signaling. Experimental Cell Research, 358(1), 31–38. 10.1016/j.yexcr.2017.06.003

10. Clé, M., Barthelemy, J., Desmetz, C., Foulongne, V., Lapeyre, L., Bolloré, K., Tuaillon, E., Erkilic, N., Kalatzis, V., Lecollinet, S., Beck, C., Pirot, N., Glasson, Y., Gosselet, F., Alvarez Martinez, M. T., Van De Perre, P., Salinas, S., & Simonin, Y. (2020). Study of Usutu virus neuropathogenicity in mice and human cellular models. PLOS Neglected Tropical Diseases, 14(4), e0008223. 10.1371/journal.pntd.0008223

11. Clé, M., Constant, O., Barthelemy, J., Desmetz, C., Martin, M. F., Lapeyre, L., Cadar, D., Savini, G., Teodori, L., Monaco, F., Schmidt-Chanasit, J., Saiz, J.-C., Gonzales, G., Lecollinet, S., Beck, C., Gosselet, F., Van De Perre, P., Foulongne, V., Salinas, S., & Simonin, Y. (2021). Differential neurovirulence of Usutu virus lineages in mice and neuronal cells. Journal of Neuroinflammation, 18(1), 11. 10.1186/s12974-020-02060-4

12. Colaneri, M., Lissandrin, R., Calia, M., Bassoli, C., Seminari, E., Pavesi, A., Rovida, F., Baldanti, F., Muzzi, A., Chichino, G., Regazzetti, A., Grecchi, C., Pan, A., Lupi, M., Franceschini, E., Mussini, C., & Bruno, R. (2023). The WEST Study: A Retrospective and Multicentric Study on the Impact of Steroid Therapy in West Nile Encephalitis. Open Forum Infectious Diseases, 10(3), ofad092. 10.1093/ofid/ofad092

13. Constant, O., Maarifi, G., Barthelemy, J., Martin, M.-F., Tinto, B., Savini, G., Van De Perre, P., Nisole, S., Simonin, Y., & Salinas, S. (2023). Differential effects of Usutu and West Nile viruses on neuroinflammation, immune cell recruitment and blood–brain barrier integrity. Emerging Microbes & Infections, 12(1), 2156815. 10.1080/22221751.2022.2156815

14. De Vries, L., & Harding, A. T. (2023). Mechanisms of Neuroinvasion and Neuropathogenesis by Pathologic Flaviviruses. Viruses, 15(2), 261. 10.3390/v15020261

15. Ershov, D., Phan, M.-S., Pylvänäinen, J. W., Rigaud, S. U., Le Blanc, L., Charles-Orszag, A., Conway, J. R. W., Laine, R. F., Roy, N. H., Bonazzi, D., Duménil, G., Jacquemet, G., & Tinevez, J.-Y. (2022). TrackMate 7: Integrating state-of-the-art segmentation algorithms into tracking pipelines. Nature Methods, 19(7), 829–832. 10.1038/s41592-022-01507-1

16. Garber, C., Soung, A., Vollmer, L. L., Kanmogne, M., Last, A., Brown, J., & Klein, R. S. (2019). T cells promote microglia-mediated synaptic elimination and cognitive dysfunction during recovery from neuropathogenic flaviviruses. Nature Neuroscience, 22(8), 1276–1288. 10.1038/s41593-019-0427-y

17. Garrido-Mesa, N., Zarzuelo, A., & Gálvez, J. (2013). Minocycline: Far beyond an antibiotic. British Journal of Pharmacology, 169(2), 337–352. 10.1111/bph.12139

18. Getts, D. R., Terry, R. L., Getts, M. T., Deffrasnes, C., Müller, M., Van Vreden, C., Ashhurst, T. M., Chami, B., McCarthy, D., Wu, H., Ma, J., Martin, A., Shae, L. D., Witting, P., Kansas, G. S., Kühn, J., Hafezi, W., Campbell, I. L., Reilly, D., … King, N. J. C. (2014). Therapeutic Inflammatory Monocyte Modulation Using Immune-Modifying Microparticles. Science Translational Medicine, 6(219). 10.1126/scitranslmed.3007563

19. Getts, D. R., Terry, R. L., Getts, M. T., Müller, M., Rana, S., Deffrasnes, C., Ashhurst, T. M., Radford, J., Hofer, M., Thomas, S., Campbell, I. L., & King, N. J. (2012). Targeted blockade in lethal West Nile virus encephalitis indicates a crucial role for very late antigen (VLA)-4-dependent recruitment of nitric oxide-producing macrophages. Journal of Neuroinflammation, 9(1), 246. 10.1186/1742-2094-9-246

20. Getts, D. R., Terry, R. L., Getts, M. T., Müller, M., Rana, S., Shrestha, B., Radford, J., Van Rooijen, N., Campbell, I. L., & King, N. J. C. (2008). Ly6c+ “inflammatory monocytes” are microglial precursors recruited in a pathogenic manner in West Nile virus encephalitis. The Journal of Experimental Medicine, 205(10), 2319–2337. 10.1084/jem.20080421

21. Glass, W. G., Lim, J. K., Cholera, R., Pletnev, A. G., Gao, J.-L., & Murphy, P. M. (2005). Chemokine receptor CCR5 promotes leukocyte trafficking to the brain and survival in West Nile virus infection. The Journal of Experimental Medicine, 202(8), 1087–1098. 10.1084/jem.20042530

22. John, C. C., Carabin, H., Montano, S. M., Bangirana, P., Zunt, J. R., & Peterson, P. K. (2015). Global research priorities for infections that affect the nervous system. Nature, 527(7578), S178–S186. 10.1038/nature16033

23. Kal, S., Beland, A., & Hasan, M. (2022). West Nile Neuroinvasive Disease Treated With High-Dose Corticosteroids. Cureus. 10.7759/cureus.31971

24. Kann, O., Almouhanna, F., & Chausse, B. (2022). Interferon γ: A master cytokine in microglia-mediated neural network dysfunction and neurodegeneration. Trends in Neurosciences, 45(12), 913–927. 10.1016/j.tins.2022.10.007

25. Käufer, C., Chhatbar, C., Bröer, S., Waltl, I., Ghita, L., Gerhauser, I., Kalinke, U., & Löscher, W. (2018). Chemokine receptors CCR2 and CX3CR1 regulate viral encephalitis-induced hippocampal damage but not seizures. Proceedings of the National Academy of Sciences, 115(38). 10.1073/pnas.1806754115

26. Kempermann, G., & Neumann, H. (2003). Microglia: The Enemy Within? Science, 302(5651), 1689–1690. 10.1126/science.1092864

27. Klein, R. S., Garber, C., Funk, K. E., Salimi, H., Soung, A., Kanmogne, M., Manivasagam, S., Agner, S., & Cain, M. (2019). Neuroinflammation During RNA Viral Infections. Annual Review of Immunology, 37(1), 73–95. 10.1146/annurev-immunol-042718-041417

28. Klein, R. S., Garber, C., & Howard, N. (2017). Infectious immunity in the central nervous system and brain function. Nature Immunology, 18(2), 132–141. 10.1038/ni.3656

29. Lee, J.-W., Mizuno, K., Watanabe, H., Lee, I.-H., Tsumita, T., Hida, K., Yawaka, Y., Kitagawa, Y., Hasebe, A., Iimura, T., & Kong, S. W. (2024). Enhanced phagocytosis associated with multinucleated microglia via Pyk2 inhibition in an acute β-amyloid infusion model. Journal of Neuroinflammation, 21(1), 196. 10.1186/s12974-024-03192-7

30. Leis, A. A., & Sinclair, D. J. (2019). Lazarus Effect of High Dose Corticosteroids in a Patient With West Nile Virus Encephalitis: A Coincidence or a Clue? Frontiers in Medicine, 6, 81. 10.3389/fmed.2019.00081

31. Lim, J. K., Obara, C. J., Rivollier, A., Pletnev, A. G., Kelsall, B. L., & Murphy, P. M. (2011). Chemokine Receptor Ccr2 Is Critical for Monocyte Accumulation and Survival in West Nile Virus Encephalitis. The Journal of Immunology, 186(1), 471–478. 10.4049/jimmunol.1003003

32. Lin, A. A., Tripathi, P. K., Sholl, A., Jordan, M. B., & Hildeman, D. A. (2009). Gamma Interferon Signaling in Macrophage Lineage Cells Regulates Central Nervous System Inflammation and Chemokine Production. Journal of Virology, 83(17), 8604–8615. 10.1128/JVI.02477-08

33. Llorente, F., García-Irazábal, A., Pérez-Ramírez, E., Cano-Gómez, C., Sarasa, M., Vázquez, A., & Jiménez-Clavero, M. Á. (2019). Influence of flavivirus co-circulation in serological diagnostics and surveillance: A model of study using West Nile, Usutu and Bagaza viruses. Transboundary and Emerging Diseases, 66(5), 2100–2106. 10.1111/tbed.13262

34. Luther, S. A., & Cyster, J. G. (2001). Chemokines as regulators of T cell differentiation. Nature Immunology, 2(2), 102–107. 10.1038/84205

35. Malemnganba, T., Rattan, A., & Prajapati, V. K. (2024). Decoding macrophage immunometabolism in human viral infection. In Advances in Protein Chemistry and Structural Biology (Vol. 140, pp. 493–523). Elsevier. 10.1016/bs.apcsb.2023.12.003

36. McGavern, D. B., & Kang, S. S. (2011). Illuminating viral infections in the nervous system. Nature Reviews Immunology, 11(5), 318–329. 10.1038/nri2971

37. Mcmanus, C. M., Liu, J. S. H., Hahn, M. T., Hua, L. L., Brosnan, C. F., Berman, J. W., & Lee, S. C. (2000). Differential induction of chemokines in human microglia by type i and ii interferons. Glia, 29(3), 273–280. 10.1002/(SICI)1098-1136(20000201)29:3<273::AID-GLIA8>3.0.CO;2-9

38. Michlmayr, D., McKimmie, C. S., Pingen, M., Haxton, B., Mansfield, K., Johnson, N., Fooks, A. R., & Graham, G. J. (2014). Defining the Chemokine Basis for Leukocyte Recruitment during Viral Encephalitis. Journal of Virology, 88(17), 9553–9567. 10.1128/JVI.03421-13

39. Morens, D. M., Folkers, G. K., & Fauci, A. S. (2004). The challenge of emerging and re-emerging infectious diseases. Nature, 430(6996), 242–249. 10.1038/nature02759

40. Nourshargh, S., & Alon, R. (2014). Leukocyte Migration into Inflamed Tissues. Immunity, 41(5), 694–707. 10.1016/j.immuni.2014.10.008

41. O’Neill, L. A. J., Kishton, R. J., & Rathmell, J. (2016). A guide to immunometabolism for immunologists. Nature Reviews Immunology, 16(9), 553–565. 10.1038/nri.2016.70

42. Pierson, T. C., & Diamond, M. S. (2020). The continued threat of emerging flaviviruses. Nature Microbiology, 5(6), 796–812. 10.1038/s41564-020-0714-0

43. Prinz, M., Jung, S., & Priller, J. (2019). Microglia Biology: One Century of Evolving Concepts. Cell, 179(2), 292–311. 10.1016/j.cell.2019.08.053

44. Purtha, W. E., Chachu, K. A., Virgin, H. W., & Diamond, M. S. (2008). Early B-Cell Activation after West Nile Virus Infection Requires Alpha/Beta Interferon but Not Antigen Receptor Signaling. Journal of Virology, 82(22), 10964–10974. 10.1128/JVI.01646-08

45. Roberts, T. K., Eugenin, E. A., Lopez, L., Romero, I. A., Weksler, B. B., Couraud, P.-O., & Berman, J. W. (2012). CCL2 disrupts the adherens junction: Implications for neuroinflammation. Laboratory Investigation, 92(8), 1213–1233. 10.1038/labinvest.2012.80

46. Rochfort, K. D., & Cummins, P. M. (2015). The blood–brain barrier endothelium: A target for pro-inflammatory cytokines. Biochemical Society Transactions, 43(4), 702–706. 10.1042/BST20140319

47. Rock, R. B., Hu, S., Deshpande, A., Munir, S., May, B. J., Baker, C. A., Peterson, P. K., & Kapur, V. (2005). Transcriptional response of human microglial cells to interferon-γ. Genes & Immunity, 6(8), 712–719. 10.1038/sj.gene.6364246

48. Russell, D. G., Huang, L., & VanderVen, B. C. (2019). Immunometabolism at the interface between macrophages and pathogens. Nature Reviews Immunology, 19(5), 291–304. 10.1038/s41577-019-0124-9

49. Simonin, Y. (2024). Circulation of West Nile Virus and Usutu Virus in Europe: Overview and Challenges. Viruses, 16(4), 599. 10.3390/v16040599

50. Singh, S., Singh, G., Tiwari, S., & Kumar, A. (2020). CCR2 Inhibition Reduces Neurotoxic Microglia Activation Phenotype After Japanese Encephalitis Viral Infection. Frontiers in Cellular Neuroscience, 14, 230. 10.3389/fncel.2020.00230

51. Sips, G. J., Wilschut, J., & Smit, J. M. (2012). Neuroinvasive flavivirus infections. Reviews in Medical Virology, 22(2), 69–87. 10.1002/rmv.712

52. Somebang, K., Rudolph, J., Imhof, I., Li, L., Niemi, E. C., Shigenaga, J., Tran, H., Gill, T. M., Lo, I., Zabel, B. A., Schmajuk, G., Wipke, B. T., Gyoneva, S., Jandreski, L., Craft, M., Benedetto, G., Plowey, E. D., Charo, I., Campbell, J., … Hsieh, C. L. (2021). CCR2 deficiency alters activation of microglia subsets in traumatic brain injury. Cell Reports, 36(12), 109727. 10.1016/j.celrep.2021.109727

53. Soto, A., De Coninck, L., Devlies, A.-S., Van De Wiele, C., Rosales Rosas, A. L., Wang, L., Matthijnssens, J., & Delang, L. (2023). Belgian Culex pipiens pipiens are competent vectors for West Nile virus while Culex modestus are competent vectors for Usutu virus. PLoS Neglected Tropical Diseases, 17(9), e0011649. 10.1371/journal.pntd.0011649

54. Spiteri, A. G., Wishart, C. L., Pamphlett, R., Locatelli, G., & King, N. J. C. (2022). Microglia and monocytes in inflammatory CNS disease: Integrating phenotype and function. Acta Neuropathologica, 143(2), 179–224. 10.1007/s00401-021-02384-2

55. Stamatovic, S. M., Keep, R. F., Kunkel, S. L., & Andjelkovic, A. V. (2003). Potential role of MCP-1 in endothelial cell tight junction ‘opening’: Signaling via Rho and Rho kinase. Journal of Cell Science, 116(Pt 22), 4615–4628. 10.1242/jcs.00755

56. Stamatovic, S. M., Shakui, P., Keep, R. F., Moore, B. B., Kunkel, S. L., Van Rooijen, N., & Andjelkovic, A. V. (2005). Monocyte chemoattractant protein-1 regulation of blood-brain barrier permeability. Journal of Cerebral Blood Flow and Metabolism: Official Journal of the International Society of Cerebral Blood Flow and Metabolism, 25(5), 593–606. 10.1038/sj.jcbfm.9600055

57. Terry, R. L., Getts, D. R., Deffrasnes, C., Van Vreden, C., Campbell, I. L., & King, N. J. (2012). Inflammatory monocytes and the pathogenesis of viral encephalitis. Journal of Neuroinflammation, 9(1), 270. 10.1186/1742-2094-9-270

58. Tinto, B., Kaboré, D. P. A., Kagoné, T. S., Constant, O., Barthelemy, J., Kiba-Koumaré, A., Van De Perre, P., Dabiré, R. K., Baldet, T., Gutierrez, S., Gil, P., Kania, D., & Simonin, Y. (2022). Screening of Circulation of Usutu and West Nile Viruses: A One Health Approach in Humans, Domestic Animals and Mosquitoes in Burkina Faso, West Africa. Microorganisms, 10(10), 2016. 10.3390/microorganisms10102016

59. Török, O., Schreiner, B., Schaffenrath, J., Tsai, H.-C., Maheshwari, U., Stifter, S. A., Welsh, C., Amorim, A., Sridhar, S., Utz, S. G., Mildenberger, W., Nassiri, S., Delorenzi, M., Aguzzi, A., Han, M. H., Greter, M., Becher, B., & Keller, A. (2021). Pericytes regulate vascular immune homeostasis in the CNS. Proceedings of the National Academy of Sciences, 118(10), e2016587118. 10.1073/pnas.2016587118

60. Tyler, K. L. (2018). Acute Viral Encephalitis. New England Journal of Medicine, 379(6), 557–566. 10.1056/NEJMra1708714

61. Vandooren, J., Geurts, N., Martens, E., Van Den Steen, P. E., & Opdenakker, G. (2013). Zymography methods for visualizing hydrolytic enzymes. Nature Methods, 10(3), 211–220. 10.1038/nmeth.2371

62. Vandooren, J., Van Damme, J., & Opdenakker, G. (2014). On the Structure and functions of gelatinase B/Matrix metalloproteinase-9 in neuroinflammation. In Progress in Brain Research (Vol. 214, pp. 193–206). Elsevier. 10.1016/B978-0-444-63486-3.00009-8

63. Vane, J. R., & Botting, R. M. (2003). The mechanism of action of aspirin. Thrombosis Research, 110(5-6), 255–258. 10.1016/S0049-3848(03)00379-7

64. Verma, S., Kumar, M., Gurjav, U., Lum, S., & Nerurkar, V. R. (2010). Reversal of West Nile virus-induced blood–brain barrier disruption and tight junction proteins degradation by matrix metalloproteinases inhibitor. Virology, 397(1), 130–138. 10.1016/j.virol.2009.10.036

65. Vilibic-Cavlek, T., Petrovic, T., Savic, V., Barbic, L., Tabain, I., Stevanovic, V., Klobucar, A., Mrzljak, A., Ilic, M., Bogdanic, M., Benvin, I., Santini, M., Capak, K., Monaco, F., Listes, E., & Savini, G. (2020). Epidemiology of Usutu Virus: The European Scenario. Pathogens, 9(9), 699. 10.3390/pathogens9090699

66. Wald, M. E., Sieg, M., Schilling, E., Binder, M., Vahlenkamp, T. W., & Claus, C. (2022). The Interferon Response Dampens the Usutu Virus Infection-Associated Increase in Glycolysis. Frontiers in Cellular and Infection Microbiology, 12, 823181. 10.3389/fcimb.2022.823181

67. Wang, P., Dai, J., Bai, F., Kong, K.-F., Wong, S. J., Montgomery, R. R., Madri, J. A., & Fikrig, E. (2008). Matrix Metalloproteinase 9 Facilitates West Nile Virus Entry into the Brain. Journal of Virology, 82(18), 8978–8985. 10.1128/JVI.00314-08

68. Wang, Y., Jin, S., Sonobe, Y., Cheng, Y., Horiuchi, H., Parajuli, B., Kawanokuchi, J., Mizuno, T., Takeuchi, H., & Suzumura, A. (2014). Interleukin-1β Induces Blood–Brain Barrier Disruption by Downregulating Sonic Hedgehog in Astrocytes. PLoS ONE, 9(10), e110024. 10.1371/journal.pone.0110024

69. Winkelmann, E. R., Widman, D. G., Xia, J., Johnson, A. J., Van Rooijen, N., Mason, P. W., Bourne, N., & Milligan, G. N. (2014). Subcapsular sinus macrophages limit dissemination of West Nile virus particles after inoculation but are not essential for the development of West Nile virus-specific T cell responses. Virology, 450-451, 278–289. 10.1016/j.virol.2013.12.021

