## Supplementary figures and images for "A central role for CCR2 in monocyte recruitment and blood-brain barrier disruption during Usutu virus encephalitis"

### Graphical abstract

#### Slide 1
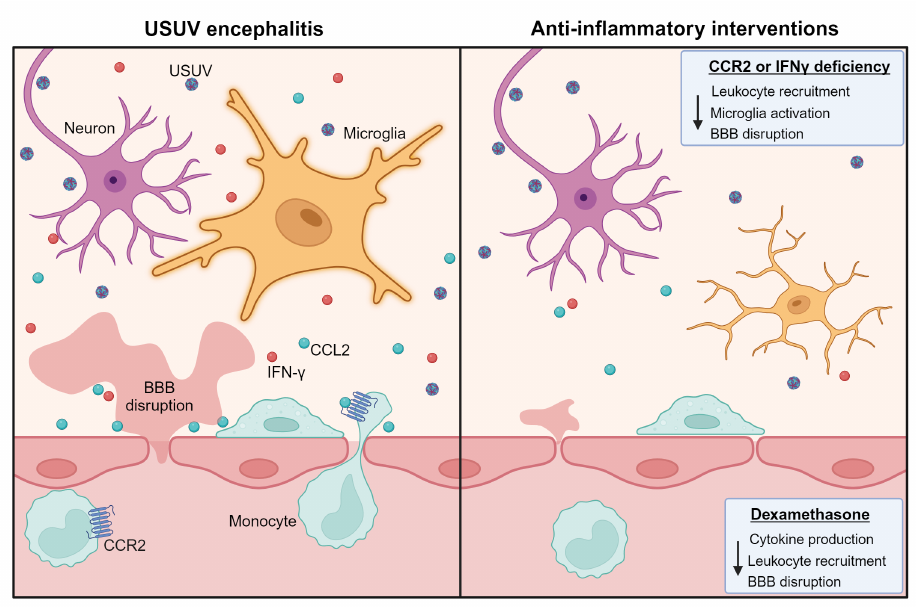
