## Supplemental data for "A central role for CCR2 in monocyte recruitment and blood-brain barrier disruption during Usutu virus encephalitis"

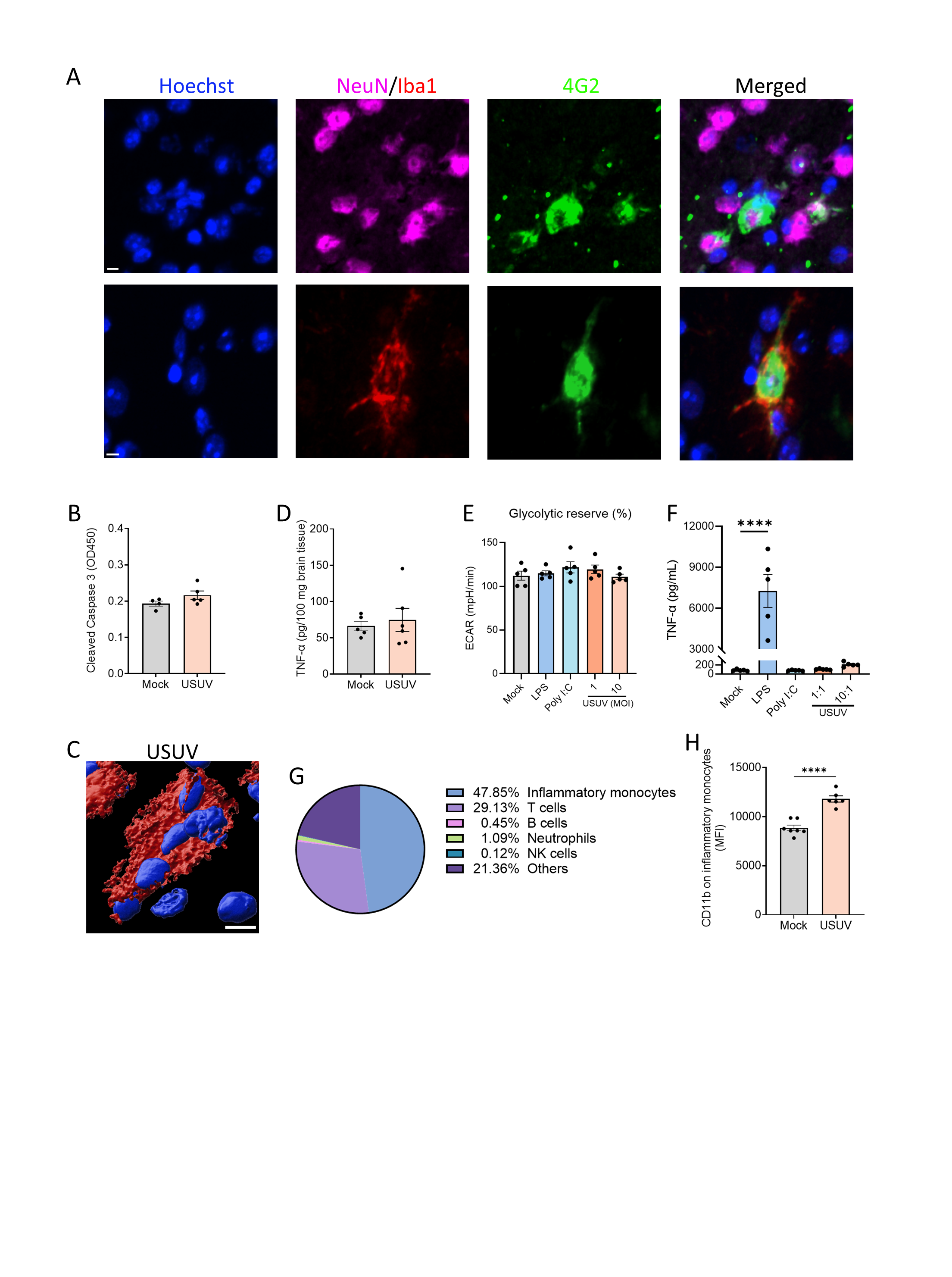

**Figure S1. Characterization of the effect of USUV infection on monocytes/macrophages. (A)** Confocal images showing immunostaining on brain cryosections from USUV-infected mice (10^4^ PFU) at 6 dpi, showing nuclei (Hoechst, blue), neurons (NeuN, magenta), microglia (Iba1, red) and USUV infection (4G2, green). Scale bars = 5 µm **(B)** Protein levels of Cleaved Caspase-3 and **(C)** Cross section of a 3D confocal image showing multinucleated microglia in brain cryosections from USUV-infected mice (10^4^ PFU) at 6 dpi. Immunostaining was performed for microglia (Iba1, red) and nuclei (Hoechst, blue). Scale bar = 10 µm. **(D)** TNF-α in brain homogenates of mock and USUV (10^4^ PFU) at 6 dpi. **(E)** Bone marrow-derived macrophages (BMDM) were stimulated with medium (mock), LPS (100 ng/mL), poly I:C (100 ng/mL) or USUV at multiplicity of infection (MOI) 1 or 10. After 24h, TNF-α levels were measured in the supernatants and **(F)** the extracellular acidification rate (ECAR) was measured from which the glycolytic reserve was calculated. **(G)** Percentages of leukocyte populations within the total group of recruited leukocytes in the brain measured by flow cytometry on 6 dpi. For gating strategy see Figure S6. **(H)** Mean fluorescence intensity (MFI) of CD11b on inflammatory monocytes in mock and USUV-infected mice, by flow cytometry. (B, D-F, H) Data are shown as mean ± SEM. N.D. = not detected. Each dot in the graphs represents a single mouse. P values were obtained with student’s t test (B, D, H) or one-way ANOVA followed by Dunnett’s multiple comparisons test (E-F). Significant differences compared to mock infected mice are indicated with *P<0.05, **P<0.01, ***P<0.001, ****P<0.0001.

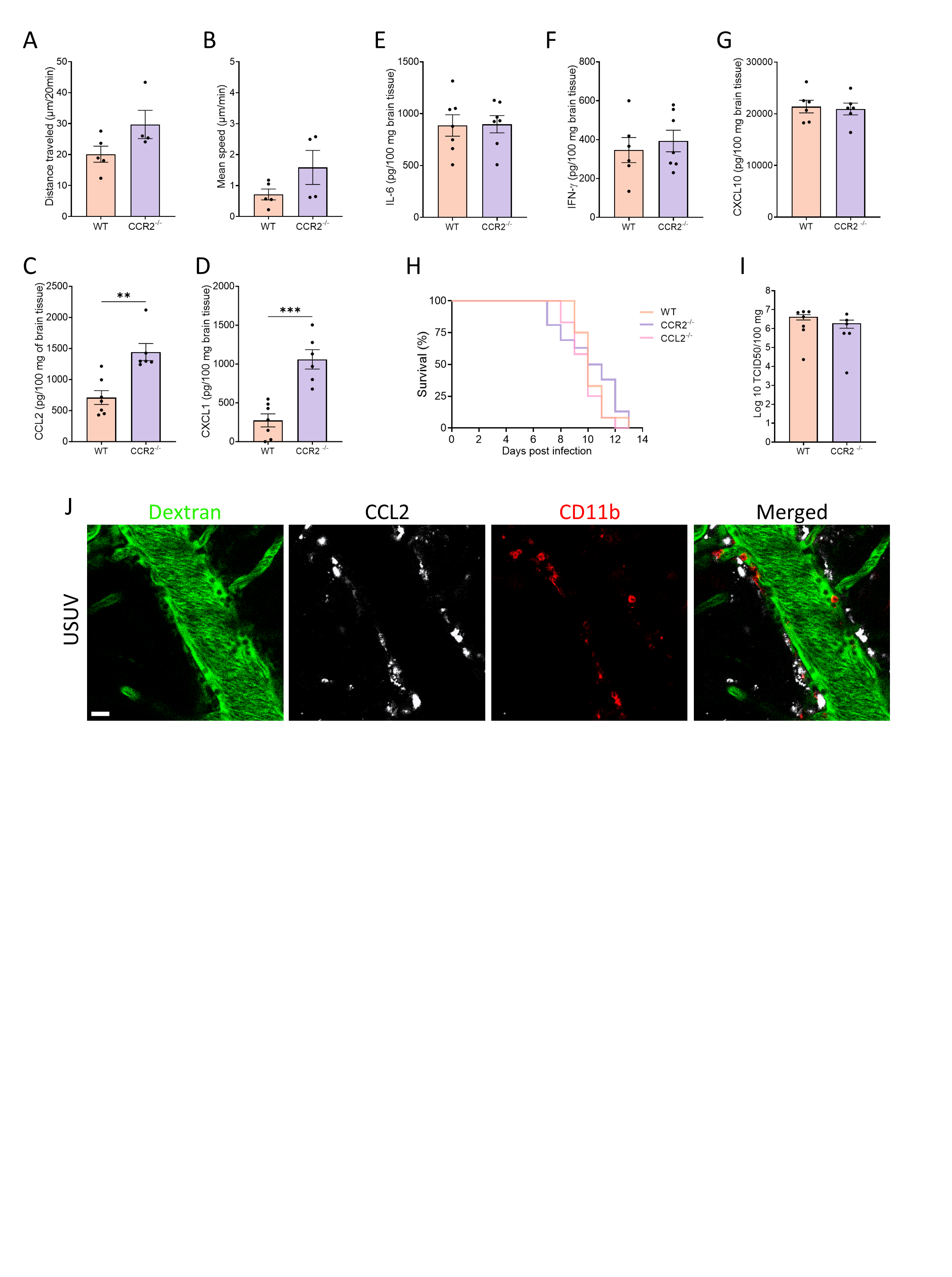

**Figure S2. CCR2 deletion does not influence the viral load or survival. (A-B)** Tracking quantification of monocytes in brain vessels of WT and CCR2^-/-^ mice from intravital microscopy videos. Tracking was performed using the plugin Trackmate 7 in FIJI. Cytokine levels of **(C)** CCL2, **(D)** CXCL1, **(E)** IL-6**, (F)** IFN-γ and **(G)** CXCL10 in brain homogenates of WT and CCR2^-/-^ USUV-infected (10^4^ PFU) mice at 6 dpi. Protein levels are displayed in pg per 100 mg of brain tissue. **(H)** Survival of WT, CCR2^-/-^ and CCL2^-/-^ mice inoculated with USUV (10^4^ PFU). N=12 per group. **(I)** Infectious virus in the brain of WT and CCR2^-/-^ mice was measured at 6 dpi and is expressed as the log_10_-transformed 50% tissue culture infectious dose (TCID_50_) per 100 mg of tissue. **(J)** Representative confocal brain intravital microscopy images from Ccl2-RFP^fl/fl^ mice at 6 dpi. Brain vasculature (dextran, green), CCL2-producing cells (RFP, white) and CD11b+ cells (CD11b, red) were visualized. Scale bar = 20 μm. Data are shown as mean ± SEM. Each dot in the graphs represents a single mouse. P values were obtained with student’s t test (A-G, I) or two-way ANOVA (H). Significant differences compared to WT infected mice are indicated with *P<0.05, **P<0.01, ***P<0.001, ****P<0.0001.

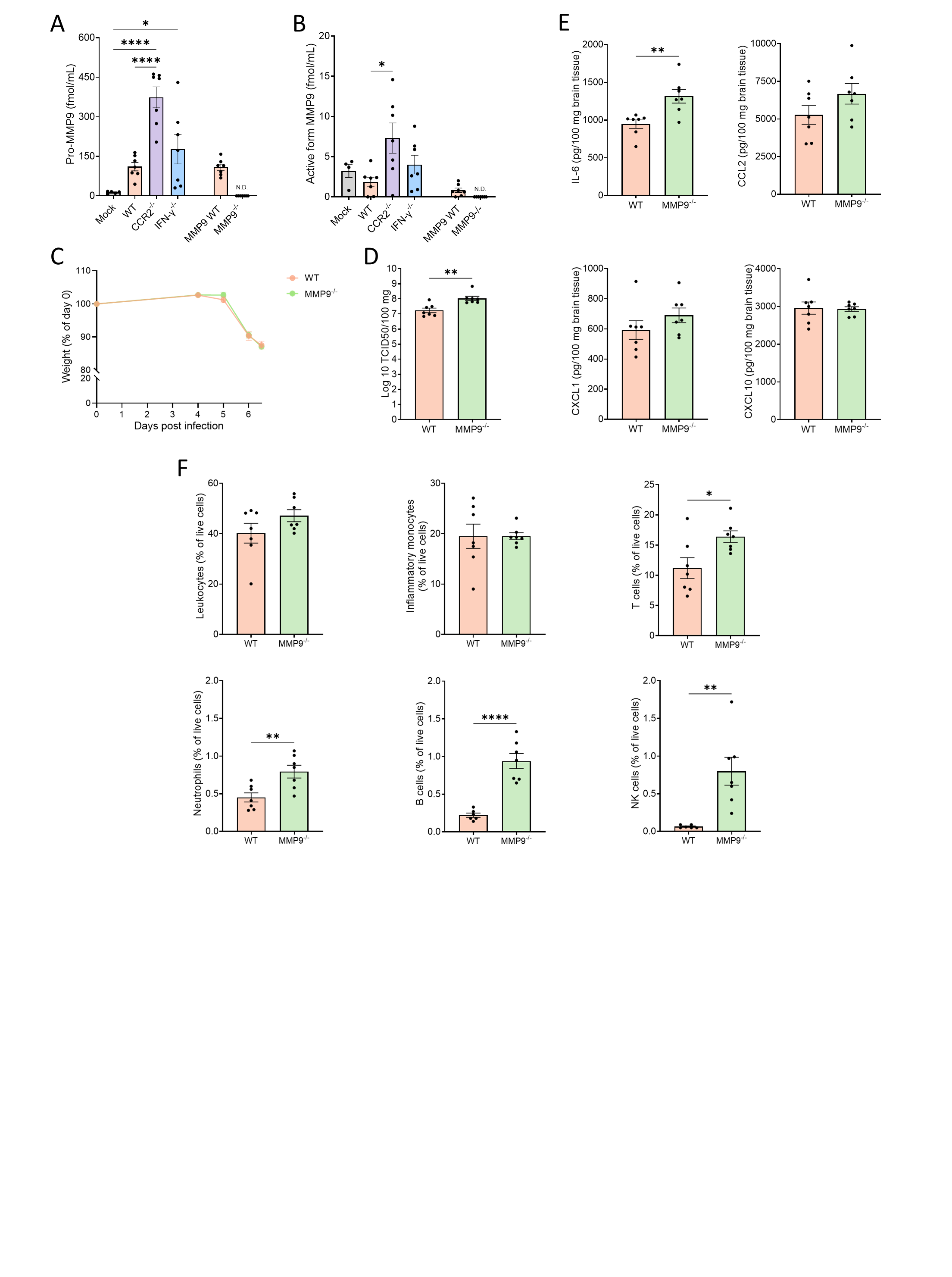

**Figure S3. MMP9 is upregulated upon USUV infection.** Quantification of gelatinase proteoforms **(A)** proMMP-9 and **(B)** the active form of MMP-9 obtained in zymographies of brain homogenates of mock (PBS), WT, CCR2^-/-^, IFN-γ^-/-^ and MMP9^-/-^ mice inoculated with USUV (10^4^ PFU). **(C)** Changes in body weight of WT and MMP9^-/-^ USUV-infected (10^4^ PFU) mice. N=7 per group. **(D)** Infectious virus in the brain of WT and MMP9^-/-^ mice was measured at 6 dpi and is expressed as the log_10_-transformed 50% tissue culture infectious dose (TCID_50_) per 100 mg of brain tissue. **(E)** Cytokine levels of IL-6, CCL2, CXCL1 and CXCL10 in brain homogenates of WT and MMP9^-/-^ USUV-infected (10^4^ PFU) mice at 6 dpi. Protein levels are displayed in pg per 100 mg of brain tissue. **(F)** Flow cytometry of cells isolated from the brain of WT and MMP9^-/-^ USUV-infected mice at 6 dpi. Percentages of total leukocytes (CD45^high^), inflammatory monocytes (CD45^high^, Ly6G^-^, CD3^-^, Ly6C^+^, CD11b^+^), T cells (CD45^high^, CD3^+^), neutrophils (CD45^high^, Ly6G^+^), B cells (CD45^high^, CD19^+^) and NK cells (CD45^high^, NK1.1^+^) within the live cell population are shown. Data are shown as mean ± SEM. N.D. = not detected. Each dot in the graphs represents a single mouse. P values were obtained with one-way ANOVA followed by Dunnett’s multiple comparisons test (A-B) two-way ANOVA (C) or student’s t test (D-F). Significant differences compared to mock (A-B) or WT USUV-infected (A-G) mice are indicated with *P<0.05, **P<0.01, ***P<0.001, ****P<0.0001.

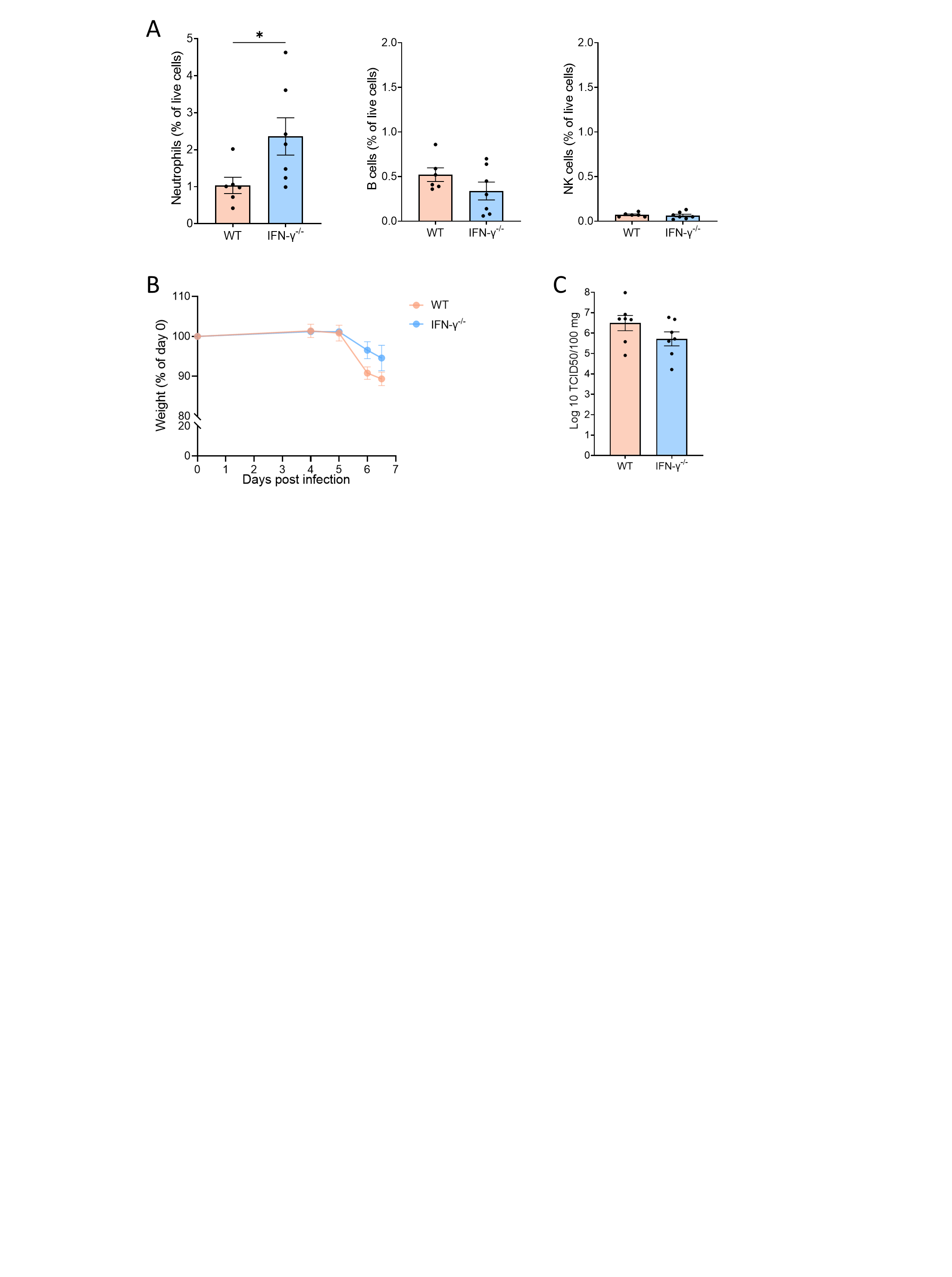

**Figure S4. IFN-γ deletion is not sufficient to protect against USUV encephalitis. (A)** Flow cytometry of cells isolated from the brain of WT and IFN-γ^-/-^ USUV-infected mice. Percentages of neutrophils (CD45^high^, Ly6G^+^), B cells (CD45^high^, CD19^+^) and NK cells (CD45^high^, NK1.1^+^) within the live cell population are shown. **(B)** Changes in body weight of WT and IFN-γ^-/-^ USUV-infected (10^4^ PFU) mice. N=7 per group. **(C)** Infectious virus in the brain of WT and IFN-γ^-/-^ mice was measured at 6 dpi and is expressed as the log_10_-transformed 50% tissue culture infectious dose (TCID_50_) per 100 mg of brain tissue. Data are shown as mean ± SEM. Each dot in the graphs represents a single mouse. P values were obtained with two-way ANOVA (B) or student’s t test (A-C). Significant differences compared to WT infected mice are indicated with *P<0.05, **P<0.01, ***P<0.001, ****P<0.0001.

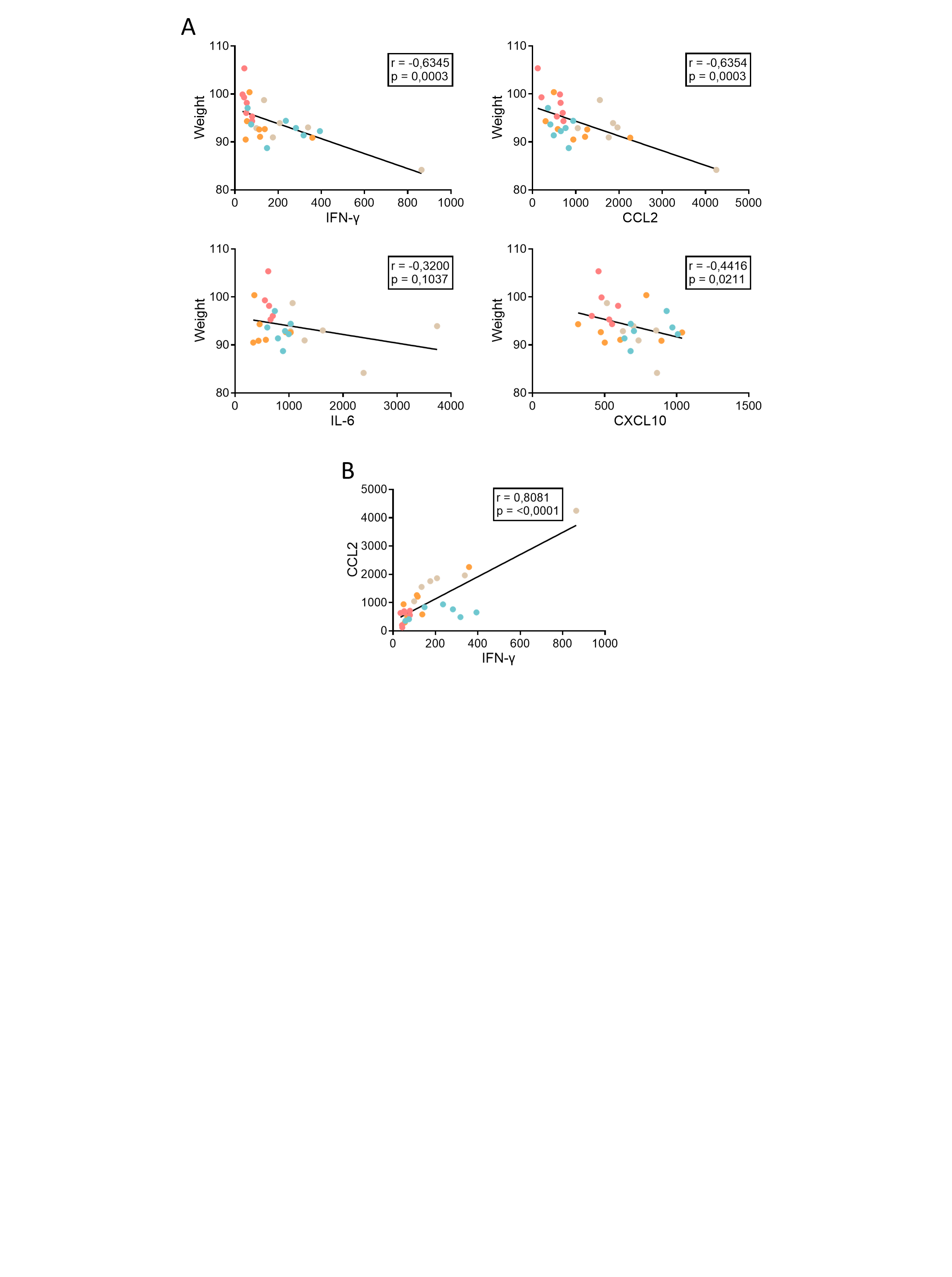

**Figure S5. Body weight and cytokine levels in the brain are negatively correlated.** C57BL/6 mice were inoculated intracranially with USUV (10^4^ PFU) and treated subcutaneously with vehicle (PBS), aspirin (100 mg/kg), dexamethasone (50 mg/kg) or minocycline (40 mg/kg) at 4, 5 and 6 dpi. **(A)** Pearson’s correlations were calculated between body weight and cytokines IFN-γ, CCL2, IL-6 and CXCL10. **(B)** Pearson’s correlation calculated between CCL2 and IFN-γ. Each dot in the graphs represents a single mouse. P values were obtained with Pearson’s correlation (A-B).

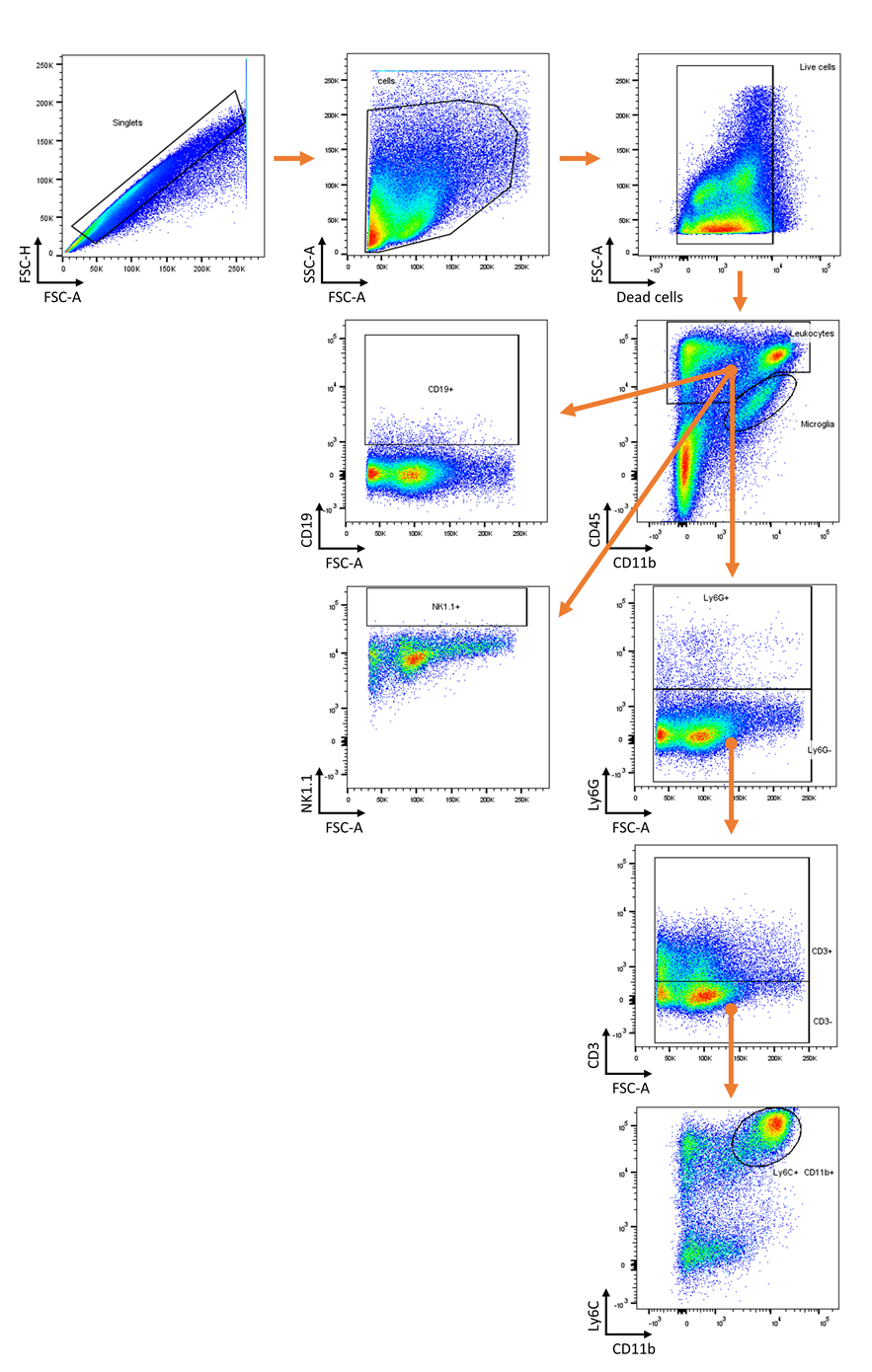

**Figure S6. Gating strategy.**

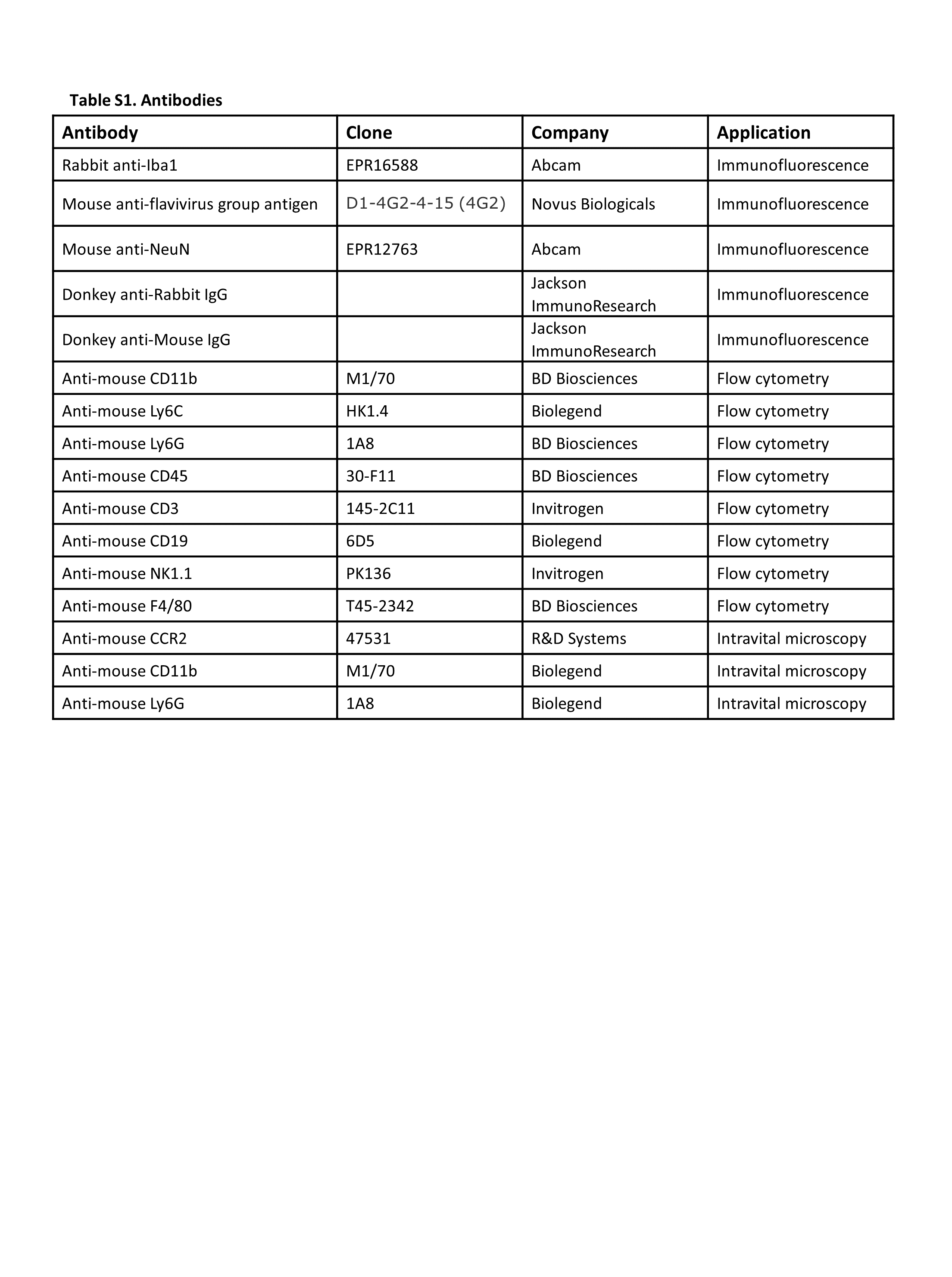
